# Regression convolutional neural network models implicate peripheral immune regulatory variants in the predisposition to Alzheimer’s disease

**DOI:** 10.1101/2022.12.02.518903

**Authors:** Easwaran Ramamurthy, Snigdha Agarwal, Noelle Toong, Irene M. Kaplow, BaDoi Phan, Andreas R. Pfenning

## Abstract

Alzheimer’s disease (AD) involves aggregation of amyloid β and tau, neuron loss, cognitive decline, and neuroinflammatory responses. Both resident microglia and peripheral immune cells have been associated with the immune component of AD. However, the relative contribution of resident and peripheral immune cell types to AD predisposition has not been thoroughly explored due to their similarity in gene expression and function. To study the effects of AD associated variants on *cis*-regulatory elements, we train convolutional neural network (CNN) regression models that link genome sequence to cell type-specific levels of open chromatin, a proxy for regulatory element activity. We then use *in silico* mutagenesis of regulatory sequences to predict the relative impact of candidate variants across these cell types. We develop and apply criteria for evaluating our models and refine our models using massively parallel reporter assay (MPRA) data. Our models identify many AD-associated variants with a greater predicted impact in peripheral cells relative to microglia or neurons but few with greater predicted impact in microglia and neurons. Our results suggest that peripheral immune cells themselves may mediate a component of AD predisposition and support their use as models to study the effects of AD associated variants. We make our library of CNN models and predictions available as a resource for the community to study immune and neurological disorders.

## INTRODUCTION

Alzheimer’s disease (AD) is a progressive age-associated neurological disorder that is characterized by neuron loss and cognitive decline. Within the brains of AD patients, aggregates of amyloid beta, aggregates of tau, and neuroinflammation can often be found well before the clinical phenotypes (1). Although the pathology of AD is neural, there is mounting evidence that AD pathogenesis is mediated in part by blood-derived cell-types, especially monocytes and macrophages. During the course of normal inflammation, monocytes from the blood infiltrate into damaged tissue and differentiate into macrophages that engulf damaged cells(2). In the case of inflammation associated with AD, monocytes cross the blood/brain barrier, differentiate into macrophages and strongly impact several molecular features of AD progression(3). These molecular functions include features that could slow AD progression, such as the clearance of amyloid beta(3, 4). However, there are also well documented neurotoxic effects that could speed up disease progression, including the induction of pathways that increase oxidative stress and secretion of proinflammatory cytokines(5). The biological model is further complicated by the fact that these functions of peripheral immune cells from the blood are also performed by resident microglia, immune cells derived from a distinct lineage(6). The relative roles of neurons, microglia and peripheral immune cells in AD predisposition and progression is still largely unknown.

Genome-wide association studies for AD have the potential to link AD predisposition to the different components of its pathology. Thus far, these studies have identified 30-40 known risk loci for Alzheimer’s disease, with hundreds more implicated at lower confidence (7–9). Despite this wealth of information, identifying candidate functional AD-associated SNPs (FAD-SNPs) remains a challenge. First, linkage disequilibrium (LD) can lead to many (up to hundreds) genetic variants at a particular locus being highly correlated. Usually, the most strongly associated variants (or sentinel variants) at disease associated loci are strong candidates for being causal variants, but it is often impossible to definitively identify which variant(s) in a region are responsible for the association signal(10). The second challenge for interpreting GWAS for AD and other studies is functional interpretation. For example, in a recent AD GWAS(11), only 2% of variants in genome-wide significant loci (p<5×10^-8^) are within known exon annotations. Instead, a large proportion of disease-associated variants lay in putative promoter and enhancer gene regulatory elements. In fact, 69% lay in enhancer states, 26% lay in regions containing promoter-associated histone marks, and 46% lay in DNase I hypersensitive sites from a large set of cell types and tissues in the HaploReg database(11, 12). Therefore, it is likely that FAD-SNPs affect the transcription of nearby genes through the alteration of regulatory element activity. Usually, this occurs by disruption of DNA sequence patterns (or motifs), which enable transcription factor (TF) binding to DNA(13). Sites of TF binding in the genome and their regulatory activities can be measured using technologies such as chromatin immunoprecipitation and sequencing (ChIP-seq) for transcription factors and histone modifications, and chromatin accessibility assays such as DNase I or transposase accessible site sequencing (DNase-seq/ATAC-seq)(14). Notably, these distal regulatory elements are highly cell type- and tissue-specific, making them a useful tool in linking the genetic loci to function (15).

Comparisons of AD GWAS to cell type- and tissue-specific epigenomic studies have highlighted the importance of the immune system in AD predisposition. Multiple studies on a variety of GWAS and regulatory datasets have found that enhancers in certain peripheral immune cells, including monocytes, are more enriched for AD-associated genetic variants than brain tissue (16, 17). In parallel, enrichments that are just as strong have been discovered in isolated and cultured microglia (17–20). For specific loci, these annotations have helped to identify the most likely function of SNPs within the locus(17, 20, 21). However, the developmental and functional similarities between peripheral monocytes and macrophages and resident microglia have made systematically dissecting the relative contributions of these cell types using standard methods difficult (22).

Several experimental methods have attempted to assign regulatory function to genetic variants. Expression quantitative trait loci (eQTL) studies have attempted to address this by identifying genetic variants associated with transcription levels of genes, but these suffer from the same LD confounds that GWAS does, so, in many cases, one cannot determine which of multiple tightly linked regulatory elements affect gene expression. In the case of AD, eQTLs have been used to identify the function of candidate loci (23, 24) and have further implicated peripheral immune cell subtypes (25), but the lack of available data in microglia makes a direct comparison difficult. Profiling of regulatory elements and careful assessment of variant effects on measured regulatory elements also could help identify causal variants. However, most profiling of regulatory elements has been conducted in a small number of non-diseased individuals, and profiling regulatory elements in large numbers of individuals with a disease of interest is usually infeasible due to high cost and challenges in collecting post-mortem tissue. This infeasibility leads to low variety of cell types and low statistical power in chromatin accessibility quantitative trait loci (caQTL)(26) and histone acetylation QTL (haQTL)(27) studies.

Machine learning-based approaches are emerging as a feasible technique to fine-map and interpret candidate regulatory disease-associated loci without the need for generating large datasets in specific cells. Building upon the availability of regulatory genomics datasets in non-diseased individuals, recent studies have developed machine learning models including convolutional neural networks (CNNs) (28–30) and gapped k-mer support vector machines (SVMs)(31–33) that can accurately predict regulatory potential of an input genome sequence in a given tissue, cell type, or context. Compared to motif scanning and position weight matrix (PWM) scanning based approaches, these models, particularly CNNs, can automatically learn motifs as well as non-linear interactions between different TF binding sites without requiring manual design of input features. These models have been shown to be more accurate than linear models whose features are known motifs. Typically, these more sophisticated models rely on the inherent genetic variation across a single human reference genome to learn the code that relates sequence to regulatory activity in a tissue, cell type, or context of interest. By learning this regulatory code, these models are capable of predicting the effect of genetic variants on regulatory activity *in silico.* Further, by direct prediction of variant effect on the overlapping regulatory element, these models are capable of fine-mapping signals from GWAS in regulatory loci and identifying the cell types, tissues, and contexts in which these variants have an effect(19).

We build upon these studies to construct a framework for identifying putative causal regulatory variants for a disease and the cell types in which they are likely to affect the disease **(Fig 1)**. In the first step, we use stratified LD score regression (S-LDSC) on open chromatin measurements from large sets of epigenomic data to identify disease relevant cell types and tissues in which transcriptional regulation is likely to contribute to disease predisposition. Then, we train CNN models on regulatory annotations in high-scoring cell types that enable the prediction of the effects of genetic variants on regulatory activity. We provide guidance on how to train these models by performing a rigorous evaluation of the performance of CNN models trained on open chromatin data to predict both regulatory activity of sequences and regulatory effects of genetic variants in a large MPRA study (34). We do this for both classification models and regression models and for models trained on both DNase-seq and MPRA data. We find that we can improve our models by using transfer learning, in which we pre-train models on open chromatin datasets and then fine tune these models on MPRA sequences. Using our model predictions, we identify which of multiple variants in LD are likely to be causal as well as the regulatory elements and cell types in which these putatively causal variants influence transcriptional regulation.

**Fig 1:**
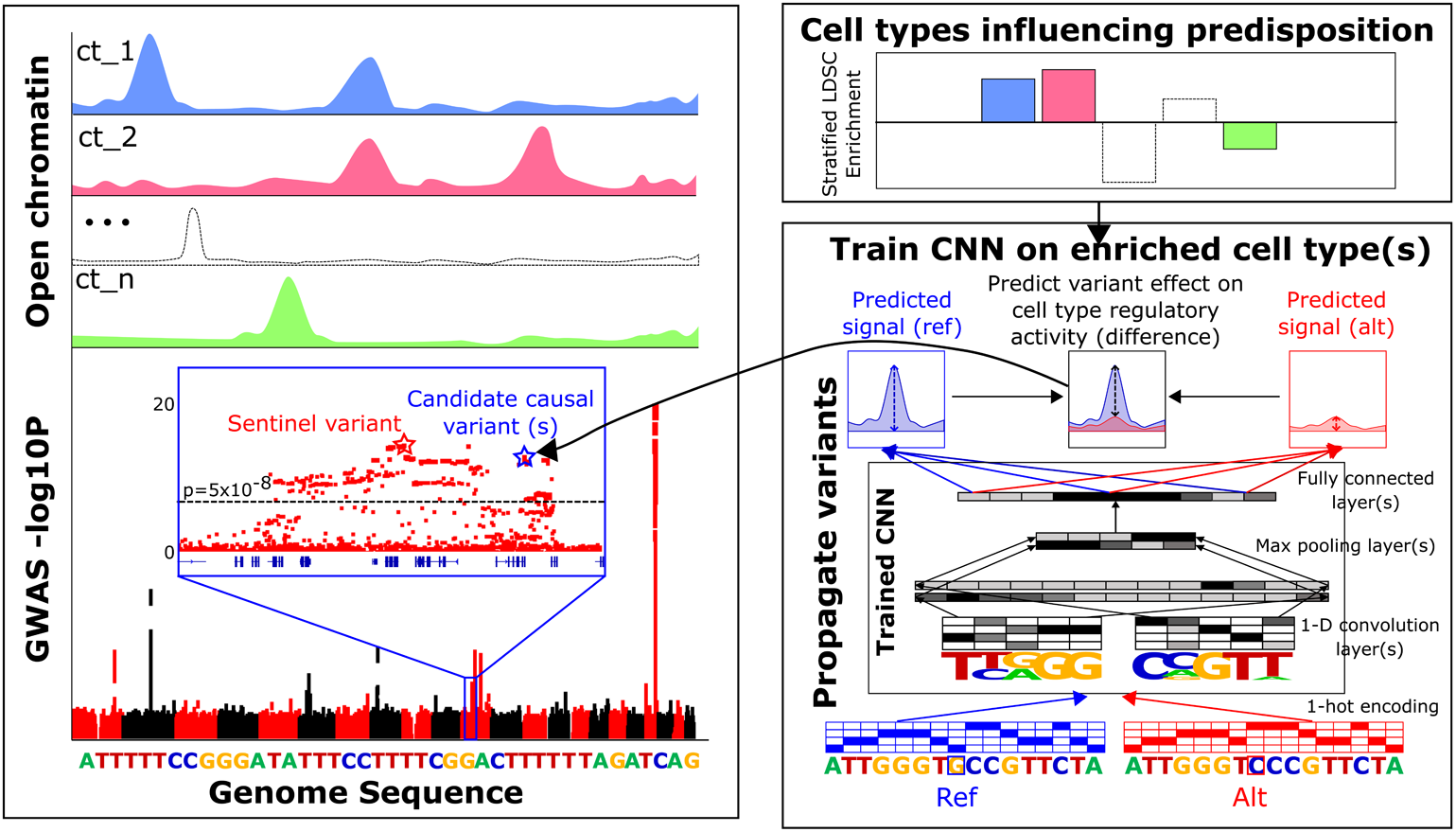
Overview of framework. In step 1, we integrate GWAS data with open chromatin profiles and use partitioned heritability analysis to identify cell types that may influence predisposition to disease through variant effects on gene regulatory elements. In step 2, we model open chromatin signal values as a function of genomic sequence using convolutional neural network (CNN) models trained on open chromatin profiles from disease relevant cell types. In step 3, we use the trained CNNs to score disease associated variants

We apply our framework to solve the dual problem of fine-mapping candidate FAD-SNPs and identifying the cell types they are most likely to act. Although previous studies have revealed that open chromatin regions in microglia are enriched for common variants associated with AD(20, 35), in a direct comparison between the two cell types using our S-LDSC analysis, we find that peripheral blood CD14+- monocyte regulatory elements are more strongly enriched for colocalization with AD associated genetic variants compared to brain resident microglial regulatory elements. We interpret the function of AD-associated genetic variants in regulatory loci that are specifically active in the peripheral blood cell types and brain resident microglia and find candidate variants that our model predicts to influence AD predisposition through disruption of regulatory activity specifically in peripheral immune cell types.

## MATERIALS AND METHODS

### Processing GM12878 open chromatin data and training CNN models on GM12878 OCRs

We reprocessed all the bulk open chromatin datasets and called irreproducible discovery rate (IDR)(36) reproducible peaks using the ENCODE ATAC-seq processing pipeline (https://github.com/ENCODE-DCC/atac-seq-pipeline), which filters low quality reads, maps reads using bowtie2(37) to the hg38 reference genome, calls peaks using MACS2(38), filters peaks in regions for which reliable read mapping may not be achievable(39), and assesses which peaks are reproducible using IDR(36). We collected 5 GM12878 DNase-seq datasets from the ENCODE data portal (Accession IDs: ENCFF057JYL, ENCFF363MQH, ENCFF716JJU, ENCFF855ODQ, ENCFF903PJT)(40, 41). Then, we obtained IDR reproducible peaks for GM12878 by treating each dataset as a biological replicate and running the pipeline on the group of datasets. We used default parameters for DNase-seq data. Since two of the replicates were deeper than the others, we used the “conservative set,” which consisted of the IDR reproducible peaks across those two replicates, as the final peak set. The conservative set is the smaller set out of the IDR reproducible peak set called across replicates and the IDR reproducible peak set called across pooled pseudo-replicates. Overall, we were able to get 110,412 peaks to train, validate, and test CNN models.

### Training regression models on GM12878 DNase-seq data

To get training data for the CNN models, we took each peak summit coordinate, expanded it to 1000 bp by extending equally in both directions and obtained the underlying sequence in the 1000 bp bins from the hg19 reference genome. To enable evaluation of model performance, we split these sequences into training, validation, and test sets by holding out all sequences derived from chromosome 4 for the validation set, and all sequences derived from chromosomes 8 and 9 for the test set. To train the CNN models, we constructed one-hot encoded arrays (1000×4 sized matrices) for the training set sequences to facilitate input into CNNs with one dimensional (1D) convolution units in the first layer. Our CNN regression models generally contained one or more convolutional layers that followed the first convolutional layer. These convolutional layers were followed by a single max pooling layer with dropout regularization(42), and then a fully connected layer with a sigmoid activation function. This sigmoid layer was followed by a single output unit. For the regression models, we used stochastic gradient descent (SGD) to minimize the mean squared error (MSE) between this output and the signal value output from the ENCODE pipeline associated with each training peak/sequence. To select the best performing architecture and hyperparameters, we evaluated model performance on the chromosome 4 held-out sequences and chose the best-performing model in terms of MSE. We fixed the number of filters in the first convolution layer to 1000 with a filter size of 8 and a stride of 1. We varied the number of convolution layers (2 or 3) following the first layer, the number of filters (filter size 8, stride 1) in these convolution layers (200 or 300), the dropout probability (0, 0.1, 0.5, or 0.8) for the max pooling layer (pooling size 13, stride=13) after the final convolution layer, the number of units in the sigmoid layer (100 or 200) that came after the max pooling layer, and the learning rate (0.1, or 0.01) and batch size (30 or 100) for SGD. Each model was trained for 100 passes through the training set (or “epochs”). We report the test set performance of the best performing model on the validation set based on mean squared error. We used Keras v2.2.4 (https://github.com/keras-team/keras) with a theano(43) backend to implement our models and train them on graphics processing units (GPUs). Since theano is now deprecated, we also provide tensorflow(44) converted model files in our data upload to facilitate future work.

### Training classifiers on GM12878 DNase-seq data

To train the CNN classifiers, we obtained negative examples of background sequences that do not underlie GM12878 OCRs. Since models of open chromatin have been shown to learn GC content profiles in sequences, we chose to use negative example sequences that matched the G+C-content of the positive example sequences. To achieve this, we used BiasAway(45, 46) to generate a set of negative sequences and associated coordinates in hg19 that matched the G+C-content distribution of our training set sequences. In order to remove false negative sequences before training, we then used bedtools subtract(47) to remove sequences in this negative set that overlapped peaks in each of the individual GM12878 DNase-seq replicates, and in the IDR optimal and conservative sets output by the ENCODE pipeline. We constructed a superset of positive and negative sequences in order to train the model. Again, we held out sequences on chromosome 4 as a validation set and sequences on chromosomes 8 and 9 as a test set. We set up the CNN classifiers similar to the regression CNN models with an initial 1D convolution layer followed by one or more additional 1D convolution layers and a max pooling layer with dropout regularization. However, the following layers differed between the classifier and the regression models: We did not use a fully connected layer with sigmoid activations for the classifier and instead, a linear transformation was applied to the outputs of the max-pooling layer which was followed by a fully connected layer and a single output unit with a sigmoid activation function which outputs a probability of whether the input sequences underlies a GM12878 OCR. We trained the model using SGD to minimize cross-entropy between this output and the binary label of peak/not-a-peak for the training sequences. Again, we selected the best performing model based on cross entropy on the chromosome 4 validation set. We varied the number of convolution layers (2 or 3) following the first layer, the number of filters (filter size 8, stride 1) in these convolution layers (200 or 300), the dropout probability (0, 0.1, 0.4, 0.5, 0.8) for the max pooling layer (pooling size 13, stride=13) after the final convolution layer, and the learning rate (0.1, or 0.01) for SGD. We used a constant batch size of 64. Each model was trained for 100 passes through the training set. We report the test set performance of the best performing model on the validation set based on mean squared error.

### Reprocessing of Tewhey et al. MPRA sequences and training CNN models on MPRA sequences

We downloaded information for all genetic variants and allelic pairs included in the 150bp oligo library with 79,000 sequences from the Tewhey et al(34) high throughput MPRA. This included information about the reference allele, alternative allele, the hg19 coordinate for each genetic variant, whether the 150bp oligo was reverse complemented, and whether the 150bp oligo library included alternative alleles for other genetic variants in the 150 bp window. Since the raw sequences were not publicly available, and we needed to construct 1000 bp sequences for propagating them through our models, we reconstructed 1000 bp sequences by putting the oligo in the center and using hg19 surrounding sequence to make up the rest of the 1000bp of sequence. In places where the original 150bp oligo was reverse-complemented, we reconstructed the reverse complement of the entire 1000bp sequence that we constructed. In addition, where usage of alternative alleles for nearby genetic variants in the 150bp were indicated, we substituted in the alternative alleles for all genetic variants in the 1000 Genomes Phase 1 data that lay within the 1000bp window. To help other researchers create machine learning model training sets using our approach, our code for reconstructing these sequences is made available at: https://github.com/pfenninglab/ml-prototypes/blob/master/lcl_regressions/construct_tewhey_sequences.py and https://github.com/pfenninglab/ml-prototypes/blob/master/lcl_regressions/parse_snp_data_for_mpra_training.py. To train models directly on MPRA sequences, we held out all sequences derived from chromosome 3 and chromosome 7 as a validation set, and all sequences derived from chromosomes 1,2,4,6,8,10,13,14,15,16,18 and 20 as a test set. We then trained a CNN regression model on the sequences derived from the remaining chromosomes to predict the final expression value [or log2FC (rna/plasmid)] of each sequence (in contrast to the DNase-seq model, which was trained to predict the signal value for open chromatin). We selected the best-performing model based on mean squared error on the validation set. We fixed the number of filters in the first convolution layer to 1000 with a filter size of 8 and a stride of 1. We varied the number of convolution layers (2 or 3) following the first layer, the number of filters (filter size 8, stride 1) in these convolution layers (200 or 300), the dropout probability (0, 0.1, 0.5, or 0.8) for the max pooling layer (pooling size 13, stride=13) after the final convolution layer, the number of units in the sigmoid layer (100 or 200) that came after the max pooling layer, and the learning rate (0.1, or 0.01) and batch size (30 or 100) for SGD. Each model was trained for 100 passes through the training set. We report the test set performance of the best performing model on the validation set based on mean squared error.

### Propagation of reprocessed MPRA sequences through LCL DNase- and MPRA-trained CNN models

For each sequence in the Tewhey et al. 79,000 oligo library, we took the 1000bp sequences constructed earlier and propagated them through the best performing CNN regression model trained on GM12878 DNase-seq, the best performing CNN classifier trained on GM12878 DNase-seq and the best-performing CNN regression model trained on the MPRA data. We then visualized the relationship between the predicted value from each of the three models and the MPRA log2FC value for each sequence in order to evaluate whether the models accurately predict regulatory activity (as measured by the log2FC value in the MPRA). To perform a quantitative evaluation of each model, we computed Pearson’s and Spearman’s correlation between predicted expression and measured expression restricting to those sequences that were included in the MPRA CNN test set.

To test whether models can predict allelic skew, we took each reference/alternative allelic pair in the 79,000 oligo library and compared model outputs for the associated pair of 1000 bp sequences. For the DNase-trained CNN regression and the MPRA-trained CNN regression model, we directly subtracted the model output for the reference allele carrying sequence from the alternative allele carrying sequence to obtain predicted allelic skew values. For the DNase-trained CNN classifier, we first performed an inverse sigmoid (or logit) transformation on the model output, then computed the difference in the transformed scores between the reference allele carrying sequence and the alternative allele carrying sequence. Genetic variants in the 79,000 oligo library were categorized into variants that displayed confident positive allelic skew (|log(skew)|≥0.2, q<0.05) and variants that did not display confident allelic skew (|log(skew)|<0.2, q>0.05). Further, variants that displayed confident allelic skew (or expression modulating variants - “emVars”) were categorized into ones that showed positive allelic skew (i.e. expression for reference allele carrying sequence was higher in the assay than the alternative allele carrying sequence) and ones that showed negative allelic skew (i.e. expression for reference allele carrying sequence was lower in the assay than the alternative allele carrying sequence). Then, we used t-tests to test whether our CNN models are able to accurately predict the direction of allelic skew. For emVars with positive skew, we tested whether the mean of predicted allelic skews from our model was greater than 0 using a one-sample one-sided t-test. For emVars with negative skew, we tested whether the mean of predicted allelic skews from our model were less than 0 using a one-sample one-sided t-test. For neutral variants that did not display a confident allelic skew in the assay, we used a one-sample two-tailed t-test to test whether the mean was significantly different from 0. We plotted the distribution of predicted allelic skews and the p-values from these t-tests for all three models in a single violin plot.

### Transfer learning

Since culturing most non-LCL non-cancer cell lines is challenging, generating large MPRAs from cells related to many diseases is difficult. In practice, only smaller MPRAs may be feasible. Therefore, we evaluated whether we could use a transfer learning approach to train a model with fewer MPRA sequences to achieve better performance at predicting enhancer activity via MPRA than a model trained on DNase-seq data alone. To do this, we trained machine learning models to predict the continuous activity of the sequences that was measured by the Tewhey *et al.* MPRA (34). We separated the MPRA sequences into training, validation, and testing sets, where the training set was derived from chromosomes 17, 11, 19, 12, 5, 22, 9, and 21; the validation set was derived from chromosomes 3 and 7; and the test set was derived from chromosomes 1, 6, 10, 4, 2, 16, 20, 15, 8, 14, 18, and 13. This gave us a training set with 38,512 sequences, a validation set with 7,627 sequences, and a test set with 33,179 sequences. We used 150 bp sequences instead of 1,000 bp padded sequences because we wanted to evaluate if our transfer learning approach could be used to create a follow-up MPRA with likely enhancer sequences, so we needed to limit our sequence lengths to those on the MPRA. We trained 3 types of models on different sized subsets of the training set. For the first model, we trained on the full training set; for the second model, we used 50% of the training set and for the third model, we used 25% of the training set. Since we used the same training/validation/test set as those used for the previous MPRA-trained regression, we started training each CNN model using the architecture from the previous model and then tuned hyper-parameters based on validation set performance, which we evaluated using mean squared error as well as Pearson and Spearman correlation coefficients between the validation set real activity and predicted activity. We trained CNN models using Keras version 2.2.4 with the theano version 1.0.3 backend, and we evaluated the models using scipy version 1.3.1. We selected the best performing model for each based on mean squared error on the full validation set. We did not downsample the data further because the validation set performance for the model trained on 25% of the dataset was not especially strong. Our architecture was similar to the earlier trained CNN regression model on 1000 bp LCL DNase peak sequences. We used an initial convolution layer with 1000 filters (filter size=8, stride=1) with a ReLU activation followed by a variable number of additional convolution layers with ReLU activations and a variable number of filters (filter size=8, stride=1) in each layer. We found that 3 additional convolution layers with 150 filters each worked best for the model trained on the full training set. For the model trained on 50% of the training set, 2 additional convolution layers with 150 filters each worked best. For the model trained on 25% of the training set, 2 additional convolution layers with 100 filters (filter size=8, stride=1) each worked best. In all models, these convolution layers were followed by a max pooling (pooling size=13, stride=13) layer. We applied dropout to the output of the max pooling layer with probability 0.8. The outputs of the max pooling layer went into a fully connected layer with a variable number of sigmoid activation units. For the models trained on 100% and 50% of the full training set, 100 units worked best for this layer, whereas for the model trained on 25% of the full training set, 50 units worked best. The output of this layer was fed to a single output unit with no activation. We found that a learning rate of 0.02 and batch size of 100 worked best for all models.

To evaluate if transfer learning could help improve the performance of models trained on 25% of the training dataset, we tried pre-training CNN regression models on sequences from DNase-seq peaks. To do this, we used IDR reproducible peaks from DNase-seq data from GM12878 from ENCODE (ENCODE Accession IDs: ENCFF057JYL, ENCFF363MQH, ENCFF716JJU, ENCFF855ODQ, ENCFF903PJT) and separated them into training, validation, and test sets using the same chromosomes for each that we used for the MPRA sequences. We added negative sequences to the DNase sequences by using genNullSeqs(32, 48) with settings genomeVersion=’hg19’, xfold = 2, repeat_match_tol = 0.02, GC_match_tol = 0.02, and length_match_tol = 0 to generate random regions of the genome with approximately the same G+C- and repetitive-element content as our peaks, and we found performance improved with these negative sequences. After hyper-parameter tuning, we found that a CNN model with 2 convolutional layers containing 200 convolutional filters (filter size=8,stride=1),followed by a fully connected layer with 100 sigmoid units and dropout 0.8, followed by a linear output, which was trained with learning rate 0.01, momentum 0, and batch size 100 for 100 epochs gave decent DNase-seq validation set performance. We then implemented transfer learning by using this trained model to pretrain a model for predicting MPRA sequences and fine-tuning that model with the same 100%, 50% and 25% of the MPRA sequences that we used in the corresponding MPRA model. We tuned the learning rate from 0.01 to 0.09 using the MPRA validation set and found that fine-tuning the model with learning rate 0.08 and another 100 epochs worked best.

### Reprocessing of bulk open chromatin datasets for microglia, CD14+-monocytes, and neurons

Using the ENCODE ATAC-seq pipeline (https://github.com/ENCODE-DCC/atac-seq-pipeline) with default ATAC-seq parameters, we processed all *ex vivo* human microglia ATAC-seq datasets from Gosselin et al(49) downloaded from the dbGaP website, under phs001373.v1.p1 (https://www.ncbi.nlm.nih.gov/projects/gap/cgi-bin/variable.cgi?study_id=phs001373.v1.p1), to generate a set of IDR reproducible human microglia OCRs. Similarly, we processed all human putamen NeuN+ ATAC-seq samples from GSE96949(50) to generate a set of neuronal OCRs using the ENCODE ATAC-seq pipeline with default parameters except for “atac.multimapping” : 0, “atac.cap_num_peak” : 300000, “atac.smooth_win” : 150, “atac.enable_idr” : true, and “atac.idr_thresh” : 0.1. We processed CD14+-monocyte DNase-seq samples (ENCODE Accession IDs: ENCFF231SIM, ENCFF000TBP, ENCFF690LNW, ENCFF000TBL) using the Kundaje Lab ATAC-Seq / DNase-Seq Pipeline with default parameters except for -enable_idr to generate a set of monocyte OCRs. For all three cell types, we used the IDR “optimal set” as the final peak set. The optimal set is the larger set out of the IDR reproducible peak set called across replicates and the IDR reproducible peak set called across pooled pseudo-replicates. Overall, we were able to get 181,122 microglia peaks, 126,727 monocyte peaks, and 177,004 neuron peaks to train, validate, and test CNN models.

### Reprocessing of Satpathy et al immune scATAC-seq data

We reprocessed the scATAC-seq dataset in Satpathy et al (51) to look for enrichments in specific immune cell types as well as train CNN regression models on them. Starting with the raw fragment files published in the study, we filtered cells of interest for ArchR(52) peak calling. We converted the fragment files to arrow files with parameters minFrags=50 to allow for a more relaxed cutoff of minimum number of fragments per cell and get more peaks per cell. These arrow files were then used to create an ArchRProject object to which cluster information was manually added based on grouping of cells into 8 cluster combinations. For each cell type, a subset project was created followed by peak calling for individual cell types. We ran this step to avoid iterative overlap peak removal across cell types which is a default peak calling functionality in ArchR. ArchR first calls peaks within each cell type using MACS2 followed by an iterative merging step to call a superset of peaks across cell types. In the iterative merging step, one peak with the highest signal may be selected and others in the vicinity might be filtered out may lead to loss of important cell type-specific peaks and peak summits. Another important part of ArchR peak calling is creation of pseudo-bulk replicates using addGroupCoverages. The parameters for this method should clearly reflect the number of cells in each cell type and the number of replicates in our dataset, so that ArchR uses all input data. If these are not correct, only a subset of the cells is used to call peaks and we may miss out on some signals. A maxReplicate value of 16 and maxCells value of 4000 was used to allow all data to be used. Peaks were then called using addReproduciblePeakSet with a q-value cutoff of 0.05 and a large enough maxPeaks setting to allow more than the default 150,000 peaks to be called. The natural log of the score generated by MACS2(38) peak calling was used as the regression label for the called peaks. Peaks on chromosomes 2,20,21,22 were moved from the training to the validation set to get larger sets for evaluating models.

### Stratified LD score regression (S-LDSC) analysis to identify AD-relevant cell types

To identify cell types enriched for AD risk, we conducted stratified LD score regression (S-LDSC)(53–55) analysis on open chromatin and H3K27ac ChIP-seq peaks derived from different cell types. We downloaded DNase-seq peaks for 53 cell types/tissues/cell lines as well as H3K27ac ChIP-seq peaks for 98 cell types/tissues/cell lines in the Epigenome Roadmap database(15). To the H3K27ac dataset, we added 12 other peak sets corresponding to 3 other cell type populations (microglia, neurons, and oligodendrocyte enriched glial) from the hippocampus and dorsolateral prefrontal cortex ChIP-seq dataset that we have previously analyzed (18). We then conducted S-LDSC analysis using the approach by the S-LDSC authors first on the DNase-seq dataset testing enrichment for each individual cell type against a background of the other 52 cell types in that dataset. We then extracted the enrichment p-values and corrected them across all 53 tests using Benjamini Hochberg FDR correction (BH FDR). We used a FDR q-value cutoff of 0.05 (q<0.05) to identify enriched cell types. We conducted two such analyses for Jansen et al(9) and Kunkle et al(8) in our analysis. Similarly, we ran S-LDSC analysis on the H3K27ac ChIP-seq dataset applying FDR correction across all 110 peak sets in the H3K27ac dataset. Since we saw enrichments for peripheral immune cell types in these primary DNase-seq and H3K27ac ChIP-seq analysis, we sought to identify which immune cell types are more strongly enriched relative to other cell types by conducting S-LDSC on the Satpathy et al immune scATAC-seq peaks. In this analysis, we ran S-LDSC analysis on peaks for each cell type against a background containing a merged set of peaks for the other 7 cell types. We applied multiple hypothesis correction across all 8 tests using BH FDR correction. To compare peripheral immune cell peaks with brain resident microglial peaks, we conducted a final S-LDSC analysis testing ENCODE CD14+ monocyte peaks against the brain resident microglial peaks from the Gosselin et al(49) study. We extracted and plotted LDSC model coefficients for these peaks sets along with uncorrected p-values for enrichment.

### Training CNN regression models on bulk open chromatin datasets for microglia, CD14+-monocytes, and neurons

We trained separate CNN regression models for each dataset, similar to the process described for the LCL DNase-seq data. To obtain negative examples for CNN classifiers, we again used BiasAway(45, 46) to generate a set of negative sequences and associated coordinates that matched the G+C-content distribution of our training set sequences. In order to remove potentially false negative sequences before training, we then used bedtools subtract(47) to remove sequences in this negative set that overlapped peaks in each of the individual replicates and in the IDR optimal and conservative sets output by the ENCODE pipeline. We constructed a superset of positive and negative sequences in order to train the model. For all models, we held out sequences on chromosome 4 as a validation set and sequences on chromosomes 8 and 9 as a test set. We used the rest of the sequences for training models. We followed the same model training strategy that we employed for the LCL DNase-seq dataset. To select the best performing architecture and hyperparameters, we evaluated model performance on the chromosome 4 held-out sequences and chose the best-performing model in terms of MSE. We fixed the number of filters in the first convolution layer to 1000 with a filter size of 8 and a stride of 1. We varied the number of convolution layers (2 or 3) following the first layer, the number of filters (filter size 8, stride 1) in these convolution layers (200 or 300), the dropout probability (0, 0.1, 0.5, or 0.8) for the max pooling layer (pooling size 13, stride=13) after the final convolution layer, the number of units in the sigmoid layer (100 or 200) that came after the max pooling layer, and the learning rate (0.1, or 0.01) and batch size (30 or 100) for SGD. Each model was trained for 100 passes through the training set (or “epochs”). We report the test set performance of the best performing model on the validation set based on mean squared error.

### Training scATAC-seq CNN regression models

We trained 8 models to predict the scATAC-seq signal in the 8 different cell types. Our models took in 500 bp of input sequences and were trained to predict the natural logarithm of the score generated by MACS2 peak calling in ArchR. Our models consisted of an initial convolution layer with 500 units (kernel size=8, stride=1) with a rectified linear unit activation (ReLu) followed by a max pooling operation (pooling size=4, stride=4). A low dropout regularization value of 0.001 when applied to the first max pooling layer yielded better performance than higher values. This was followed by two more convolution + max pooling blocks. The first of these contained 250 convolution filters (kernel size=8, stride=1) with ReLu activations followed by max pooling (pooling size=4, stride=4). The next one contained 100 convolution filters (kernel size=8, stride=1) with ReLu activations followed by max pooling (pooling size=4, stride=4). The output of the pooling operation was flattened and passed into a single output unit which predicts the scATAC-seq peak signal. We applied dropout with probability 0.25 and L2 kernel regularization of 0.01 to the last 2 max pooling layers. We used a cyclic learning rate(56) scheduler for SGD optimization to achieve faster convergence. Training models with an unweighted mean squared error loss function did not lead to satisfactory performance on peaks with higher signal values. Therefore, we used a weighted mean squared error loss with up-weighting for samples with a signal value greater than 4 by a factor of 3. Essentially, this penalized wrong predictions three times as heavily for higher signal instances which our earlier models were failing to predict as opposed to lower signal instances and helped improve model performance. To report final performance, we report Pearson and Spearman’s rank correlation coefficient between model predictions and true scATAC-seq signal values. Additionally, to test whether our models were predicting cell type-specific signals and differentiate between different blood cell types, we selected a cell type (monocyte) and evaluated predictions of the model trained on that cell type on peaks in another cell type (CD4+ T-cells) that did not overlap monocyte peaks. We then compared monocyte model predictions for these peaks with monocyte model predictions for monocyte peaks that did not overlap CD4+ T-cell peaks **(Fig 6E)** using a 2 sided Mann-Whitney test. As another way to test robustness of our scATAC-seq trained models, we compared predictions scATAC-seq model predictions of variant effects for a cell type (monocyte) with predictions of variant effect from models trained on bulk open chromatin datasets from the same cell type (previously trained ENCODE CD14+ bulk DNase-seq model) **(Fig 6F).** We computed Pearson’s and Spearman’s rank correlation between these variant effect predictions to see whether scATAC-seq model predictions agree with bulk open chromatin trained models.

### Constructing a set of LD expanded AD associated variants

To identify a set of input AD associated variants that we score for their effects using our CNN models, we curated a set of sentinel AD GWAS variants identified in Jansen et al.(9) and Kunkle et al(8) with a significance threshold of 5×10^-8^. We then expanded this list to include variants in linkage disequilibrium(LD) with the sentinel variants. using HaploReg(v4.1)(12), which uses LD information from the 1000 Genomes Project. Using a LD r^2^ threshold of 0.5, we obtained a total of 1,209 variants in the LD expanded set.

### Scoring of AD variants for their regulatory effects through CNN models in different cell types

We scored OCR overlapping AD associated variants in the LD expanded set using 11 CNN regression models. This includes the 3 bulk open chromatin trained models (Gosselin microglia, ENCODE CD14+ monocyte, Fullard Neuron) and the 8 Satpathy et al immune scATAC-seq trained models. For each variant, we constructed a pair of sequences, one carrying the reference allele and the other carrying the alternate allele. Surrounding sequence was obtained from the hg19 reference genome. To match the 1000nt input sequence length for the 3 bulk open chromatin trained models, we added 499 nts to the left of the variant position and 500 nts to the right of the variant position. For the scATAC-seq trained models, we added in 249 nts to the left of the variant position and 250 nts to the right of the variant position to get input sequences of length 500nt. We then predicted variant effect scores for all 11 models by computing the difference between the model predicted outputs for each of the 1,209 sequence pairs. To interpret and identify significant variant effect scores, we needed a null distribution to compare these scores against. To construct this null distribution, we obtained a full set of variant annotations across the entire genome from our collaborators at the Rudolph Tanzi Lab(57). For each of the 11 models, we then scored all variant effects in the Tanzi variant set that overlapped peaks in the dataset that was used to train/test/validate the model. We restrict to OCR overlapping variants since CNN models are known to have problems generalizing to areas of the input space that are far away from the training set distribution. Another reason for doing so was computational efficiency since scoring millions of variants even with one forward pass can take 2-3 days on a GPU. Our variant effect scores distribution were centered at 0 with Gaussian-like tails on both sides. The 0 centering is expected since most variants are unlikely to have any effects on regulatory activity. We then used properties of the Gaussian distribution to identify outlier variants that have significant effects on regulatory element activity. In our initial analysis, we identified outliers by selecting all OCR-overlapping variants that lie at least 2 standard deviations away from the 0 value. We use this cutoff because around 95% of values lie within 2 standard deviations of the mean of a Gaussian distribution. Roughly, this corresponds to a p-value cutoff of 0.05 which has been widely used in scientific literature ever since it was described in R.A. Fisher’s tables(58, 59). In secondary analysis, we scored all 1209 AD associated sequence pairs through all 11 models but only focused our interpretation on outlier variants identified in the initial analysis. We did so to enable comparison of model scores for the previously identified OCR overlapping outlier variants in other cell types where the variant does not overlap an OCR.

### Model interpretation using DeepSHAP

For specific outlier variants like rs636317 and rs76726049 identified by our ENCODE CD14+ monocyte model, we conducted model interpretation and assigned contribution scores to each input nucleotide using DeepSHAP(60). Internally, DeepSHAP builds on a connection with DeepLIFT(61) which describes the difference in output from a selected ‘reference’ output in terms of the difference of the input from the some ‘reference’ input. Usually, the reference input represents some neutral or default input that is selected by the user according to the task. We applied DeepSHAP separately on the reference allele carrying sequence and the alternate allele carrying sequence for a given variant to compute contribution scores for every nucleotide in the 1000nt input sequence. We used a reference input of all 0s which corresponds to an input sequence with all unknown nucleotides (“N”). We then plotted DeepSHAP scores for the middle 200nt of the input sequences to enable easy visualization of the central variant location.

## RESULTS

### CNN regression models and CNN classifiers trained on trained on LCL open chromatin data can accurately predict open chromatin signal from sequence

To investigate the types of models that work well for our framework, we trained CNN classification and regression models on open chromatin regions (OCRs) identified from 5 DNase-seq datasets of GM12878 lymphoblastoid cell lines (LCLs) in the ENCODE database(40, 41). We chose this cell line because it has publicly available high-throughput MPRA data that includes sequences with different alleles of genetic variants(34). We trained the classifiers on positive example sequences underlying OCRs and an almost equal number of negative example sequences derived from closed regions of the genome that were matched for G+C nucleotide content with the positive examples (**Fig 2A**). Our CNNs are similar to previously described CNN classifiers(28–30) and gapped k-mer (gkm) kernel SVM(31, 32, 62) methods in the literature with model binary labels representing open or closed chromatin. Our CNN classifier was able to accurately generalize to test set OCR and negative examples on chromosomes (chromosomes 8 and 9) not used for training or hyper-parameter tuning (area under receiver operating characteristic auROC=0.96, area under precision-recall curve auPRC=0.97) **(Fig 2B and 2C).**

**Fig 2:**
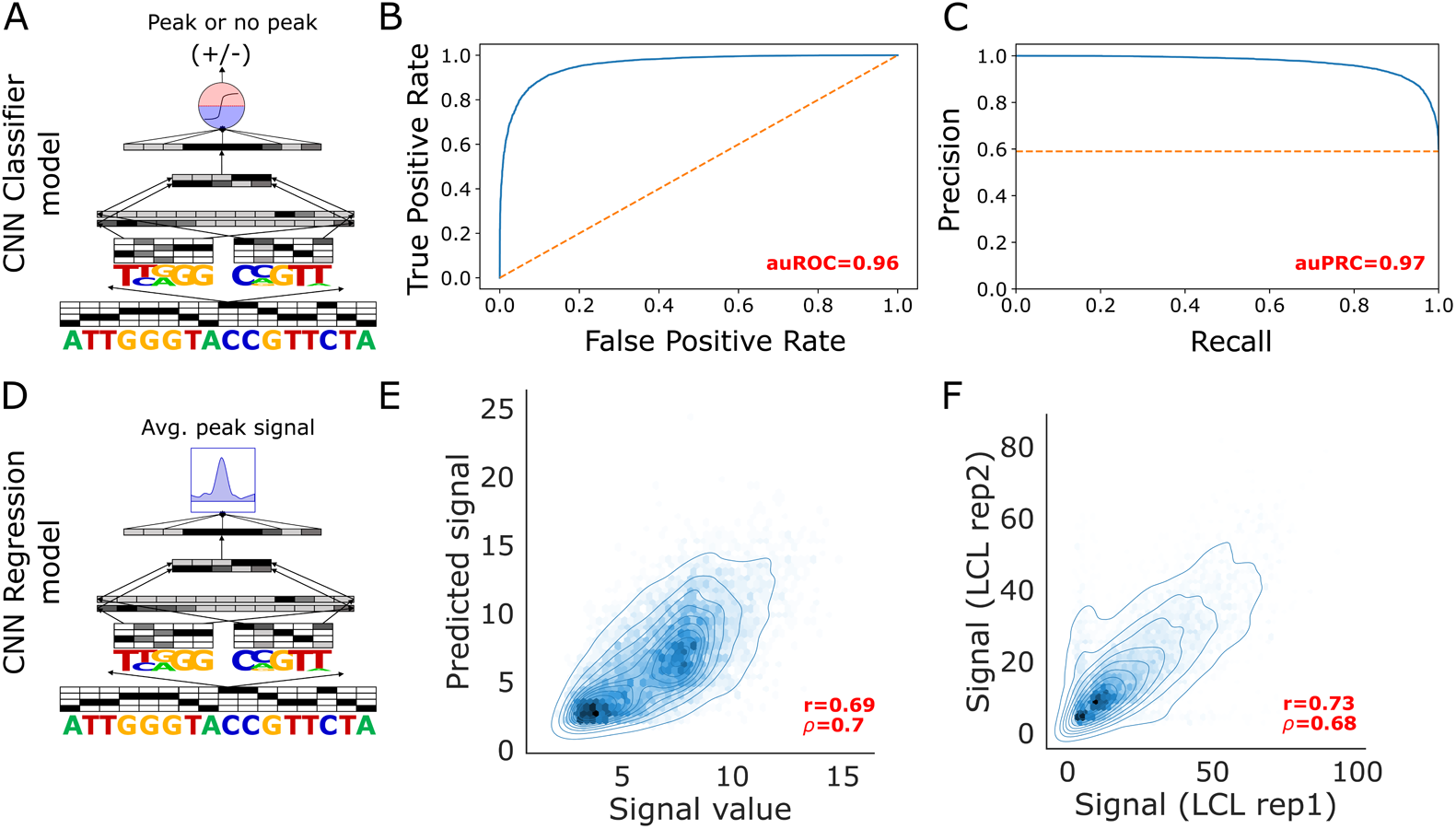
Generalization performance of CNN classifiers and regression models trained on LCL DNase-seq data. **A.** cartoon diagram of a CNN classifier that we trained. The models takes in a one-hot encoded genomic sequences, passes it through the convolutional network that contains alternating convolutional and max pooling layers and outputs a sigmoid transformed value representing the probability that a sequence underlies an OCR or not. **B.** auROC on test set sequences derived from held out chromosomes (chromosomes 8 and 9) of a classifier trained on LCL DNase-seq OCR peaks and equal numbers of GC matched negative controls **C.** auPRC of CNN classifier on held out test set sequences **D.** cartoon diagram of a CNN regression model that we trained. The model structure is similar but instead of a sigmoid probability value, it is trained to output the quantitative signal from the open chromatin assay (DNase-seq/ATAC-seq) **E.** Scatter plot of CNN regression model predicted signal vs the actual signal value from the peak calling algorithm for chromosome 8 and 9 test set peaks that were held out during training and hyperparameter tuning. Pearson and Spearman correlation between predicted and true signal is included in red text **F.** Scatter plot of true signal values in individual DNase-seq replicates for the same set of peaks presented in E

In addition, we trained CNN regressions that model quantitative OCR signal instead of a binary label **(Fig 2D)**. Our motivation for doing this is that most common variants are unlikely to fully deplete regulatory activity **(Fig S1)** and rather might have smaller effects on regulatory activity(63). Our CNN regression model was also able to generalize to held out examples on chromosome 8 and 9 LCL OCRs with a Pearson’s correlation (r) of 0.69 and Spearman’s correlation (ρ) of 0.7 between the true OCR signal and model predicted OCR signal **(Fig 2E)**. This represents a very accurate model considering that signal values for the same set of OCRs displayed a moderately higher correlation (r=0.73, ρ=0.68) between the two replicates that were used to define the final peak set that went into training **(Fig 2F)**. These results suggest that using regression models for our framework is feasible.

### LCL DNase trained CNN models can accurately predict measured regulatory activity in MPRA conducted in LCLs

Since not all OCRs function as enhancers(64), we evaluated our models on direct measurements of enhancer activity using previously published MPRA data from LCLs(34). First, we evaluated how well our models were able to predict the regulatory activity of each oligonucleotide in the assay. To conduct an unbiased evaluation, we trained another CNN that models the MPRA measured regulatory activity (log2(RNA/plasmid DNA)) as a function of the 1000bp extended sequences derived from the MPRA. This differentiates our work from previous studies that either train only on MPRA(65) or do not compare models trained on assays like DNase-seq against an MPRA-trained model(66). Our MPRA-trained model was able to generalize well with strong correlations between predicted MPRA activity and measured MPRA activity for held out test set sequences not used for training or hyperparameter tuning **(Fig 3A)**. On test set sequences that were called active in the MPRA, the MPRA-trained model achieved moderate correlation (r=0.34,ρ=0.28). Not surprisingly, the MPRA-trained model predictions displayed a lower correlation for the subset of MPRA sequences that were called inactive in the Tewhey et al. study (r=0.24, ρ=0.24). Interestingly, on the active test set sequences, our DNase trained regression model achieved moderate but statistically significant correlations between model predictions and MPRA measured signal (r=0.28,ρ=0.22) **(Fig 3B)**. We obtained slightly lower correlation values for the DNase-trained CNN classifier (r=0.25,ρ=0.22) **(Fig 3C**). Although the models trained on DNase-seq do not work as well as the MPRA-trained model, these correlations suggest that the DNase-seq-trained models are able to learn relevant signals and generalize to MPRA data to some extent.

**Fig 3:**
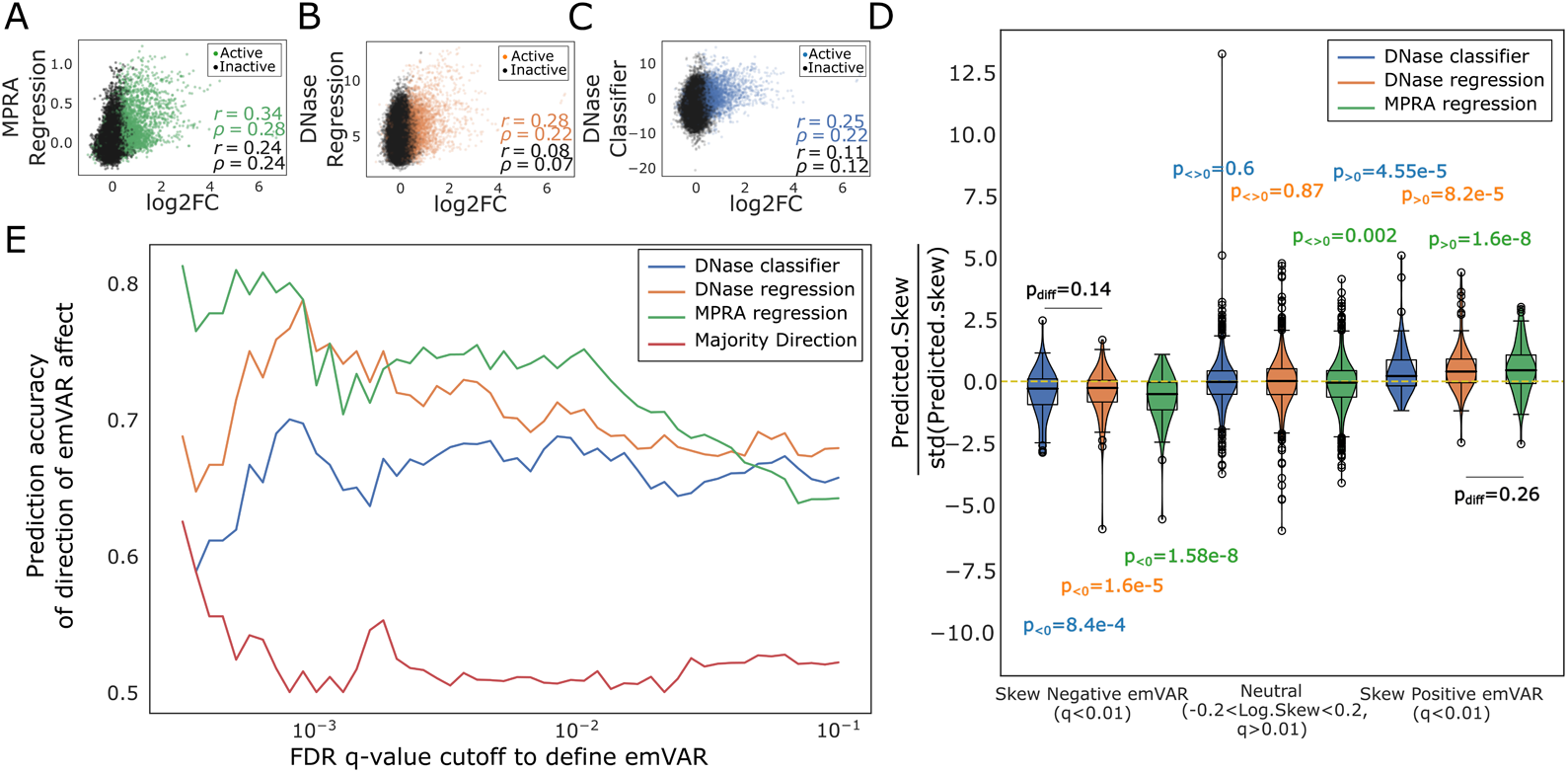
Performance of LCL DNase trained CNN classifiers, LCL DNase trained regression models and MPRA trained models on the task of predicting regulatory activity and variant effects in the Tewhey et al MPRA. **A.** Scatter plot between true and predicted MPRA regulatory activity showing generalization performance of the MPRA trained model on test sequences held out during training of the model. Pearson and Spearman correlation values between true MPRA activity and predicted MPRA activity are indicated in the bottom right for active and inactive sequences in the MPRA study **B.** Scatter plot between true MPRA signal and predicted signal from the DNase trained CNN regression model on the MPRA model test set. Points are colored by whether the sequences were deemed active or inactive in the MPRA and corresponding correlations on these subsets are also reported in the bottom right. **C.** similar to B but instead of the CNN regression, the plot displays performance and predictions from the DNase trained CNN classifier **D.** Violin plot displaying performance of the MPRA trained baseline model, the DNase trained classifiers, and the DNase trained regression on predicting allelic skew (or variant effect) in the MPRA study. Shown are the allelic skew values for three different subsets of variants derived from the MPRA test dataset. These subsets correspond to 1) variants which are confidently (FDR q<0.01) called to have higher MPRA activity for the alternate allele compared to the reference allele (i.e. ”skew negative emVars”); 2) variants that do not display confident allelic skew in the study (|log(skew)|<0.2 and FDR q>0.01); 3) variants which are confidently called to have lower MPRA activity for the alternate allele compared to the reference allele (FDR q<0.01). p-values from one-sample t-tests comparing distribution shifts from neutrality (0 value) are indicated above or below the violins. p-values from two sample t-tests (represented p_diff_) comparing predictions from the DNase trained classifier and the DNase trained regression are also displayed **E.** accuracy of the prediction of the direction of variant effect of the three models on variants in the MPRA test dataset. Accuracy is reported at different FDR q-values to call emVars. Accuracy for a naïve baseline classifier that always predicts the direction of effect for a variant to be the direction of effect for majority of variants is also included as a baseline

### Pre-training MPRA-based models with DNase-seq-trained models enables MPRA-based models to achieve good performance with less MPRA data

While our MPRA-trained models worked better than our DNase-seq trained models, our MPRA-trained models used an MPRA with over 70,000 sequences, and generating such large MPRAs for most non-cancerous cell types is infeasible due to challenges involved in culturing non-cancerous cells. Therefore, we trained models on smaller subsets of the training data to identify how many samples would be needed to train good MPRA models. As expected, we found that reducing training set size led to lower generalization performance on MPRA test set sequences **(Fig S2)**. We therefore evaluated model performances in a transfer learning framework where we pre-trained models on the DNase-seq data and fine-tuned them on a subset of the MPRA data. For pre-training, we used a CNN regression model on LCL DNase-seq defined OCRs with GC content-matched negative sequences that were labeled as having 0 signal. We found that this model generalized well to DNase-seq defined OCRs on validation set chromosomes that were not used for training (Pearson’s r=0.794 for DNase-seq defined OCRs, Spearman’s ρ=0.766). Then, we fine-tuned this DNase-seq pretrained model for MPRA prediction on the same subsets of the MPRA training dataset that were used to create the MPRA-trained models. We found that fine-tuning the model on 25% of the MPRA training dataset led to reasonable performance on held out validation set sequences (r=0.393, ρ=0.547). Strikingly, we found that the transfer learning model fine tuned on 25% of the MPRA training set performed much better at predicting a completely held out MPRA test set (r=0.394, ρ=0.381) than both the MPRA only model trained on 25% of the MPRA training set (r=0.297, ρ=0.227) and the model trained on DNase-seq without fine-tuning (r=0.304,ρ=0.22) **(Fig S2)**. This suggests that deriving information from larger datasets that assay open chromatin and fine tuning models on MPRA data can improve prediction performance for sparser MPRAs.

### CNN models trained on DNase-seq data can predict direction of variant effects in MPRA accurately

Since our final goal is to identify genetic variants that affect enhancer activity, we evaluated our models on their ability to predict the direction of allelic skew for expression modulating genetic variants (emVars). We found that the DNase-seq trained classifier and regression achieved performance on par with the MPRA trained model **(Fig 3D).** At most q-value cutoffs to define emVars, the DNase trained regression outperformed the DNase trained classifier and at some cutoffs, it achieved performance on par with the MPRA trained model **(Fig 3E)**. In addition, all three models performed markedly better than a theoretical naive classifier that predicts the direction of effect to be the majority class.

### Stratified LD score regression analysis identifies brain resident microglia and peripheral immune cell types relevant for study of regulatory variation associated with Alzheimer’s Disease

The first step of our framework involves using S-LDSC to identify cell types that are likely to be associated with disease. We apply this approach to compare the contribution of different neural and immune cell type subtypes to AD predisposition.. To do this, we first systematically conducted S-LDSC analysis to test whether heritability for AD was enriched in open chromatin and H3K27ac ChIP-seq profiles in specific cell types/tissues out of a large set of cell types/tissues. We used GWAS summary statistics from two different AD studies: Kunkle et al(8) and Jansen et al(9). From DNase-seq profiles of 53 different cell types/tissues in the Epigenome Roadmap database(15), we detected statistically significant colocalization between AD-associated variants and open chromatin in peripheral immune and blood cell types, particularly myeloid cell types. This included primary CD14+ monocytes, primary monocytes from peripheral blood, and primary natural killer (NK) cells from peripheral blood **(Fig 4A)**. Notably, microglia are not included in the Epigenome Roadmap DNase-seq datasets at all and, hence, we added microglial datasets to subsequent S-LDSC analyses. First, we conducted S-LDSC analysis on H3K27ac ChIP-seq peaks from all 98 different tissues/cell types in the Epigenome Roadmap. In addition to these 98 peak sets, we included 12 additional peak sets derived from microglia, oligodendrocyte enriched glia (OEG), and neurons from the hippocampal and cortical H3K27ac ChIP-seq dataset of non-diseased human controls that we have previously analyzed(35). In this analysis, we again found that multiple peripheral blood cell types were enriched for colocalization with AD risk **(Fig 4B)**. In addition, the microglial H3K27ac peaks displayed a significant colocalization with AD risk, in line with previous work (20, 35, 67). Further, the spleen was enriched, which is likely due to the high proportion of monocytes in that tissue(68). Notably, immune cell type enrichments on the broader H3K27ac peaks are consistent with the enrichment results on the narrower DNase-seq OCRs.

**Fig 4:**
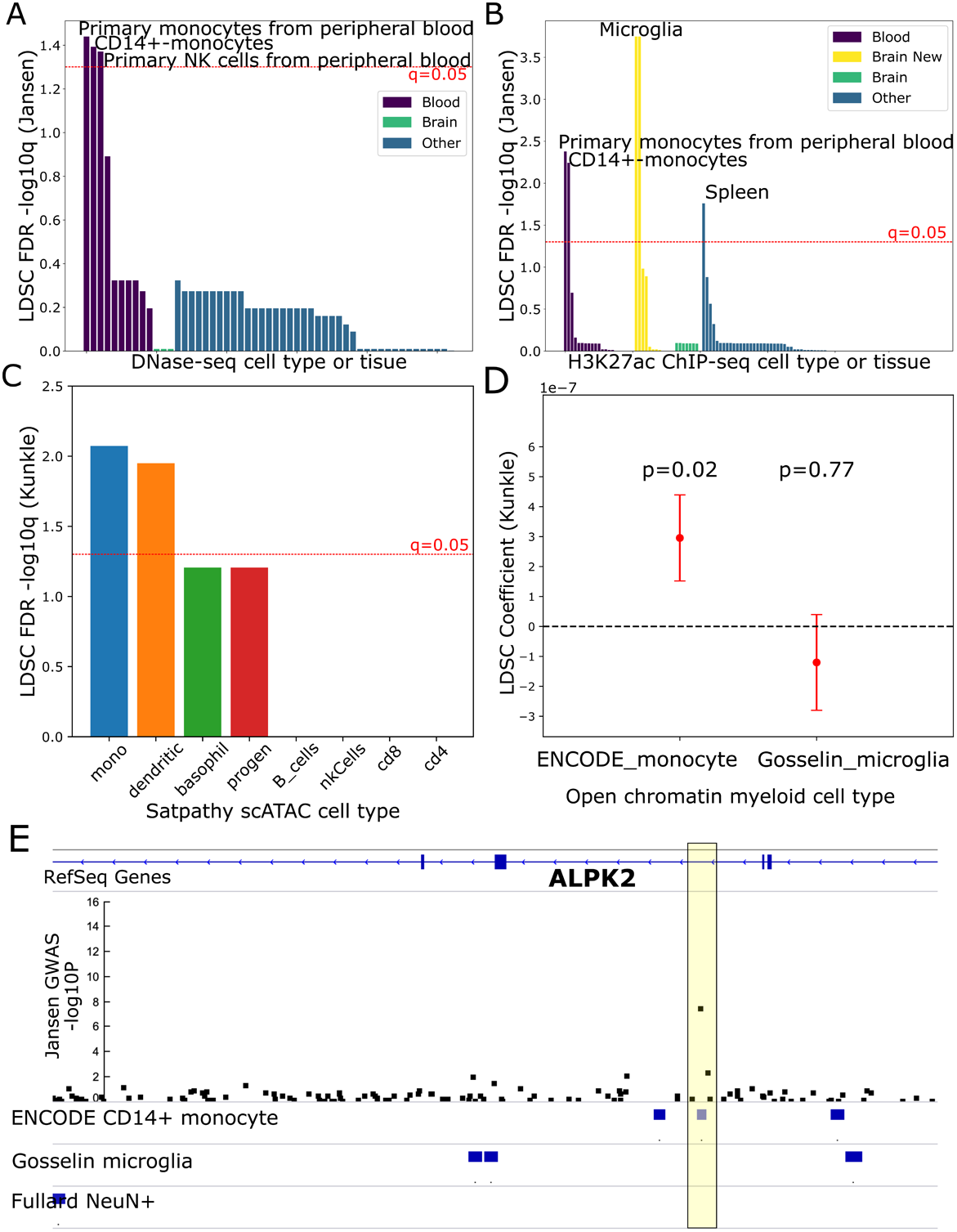
Systematic stratified LD score regression analysis on histone acetylation and open chromatin profiles reveals a stronger AD GWAS enrichment for peripheral immune cell types compared to microglia. **A.** Bar chart depicting FDR q-values from an S-LDSC analysis on the Jansen et al AD-by-proxy GWAS and included DNase-seq peaks from 53 cell types/tissues in the Roadmap Epigenomics dataset. Red line indicates q=0.05, the significance cutoff based on FDR. Tissues are categorized and colored by whether they are derived from brain, blood, or other tissues. **B.** Bar chart depicting FDR q-values from an S-LDSC analysis on the Jansen et al GWAS and H3K27ac ChIP-seq peaks from 98 cell types/tissues in Roadmap Epigenomics dataset as well as 12 profiles (”Brain New”) from the cell type-specific brain H3K27ac dataset from Ramamurthy, Welch et al. Tissues are categorized and colored by whether they are derived from brain, the new brain dataset, blood, or other tissues **C.** Bar chart depicting FDR q-values from S-LDSC analysis on the Kunkle et al GWAS and 8 immune cell type peak sets derived from the scATAC-seq dataset in Satpathy et al. mono-Monocyte, dendritic - Dendritic cells, progen - progenitors, basophil - Basophils, nkCells - Natural Killer cells, B_cells - B cells, cd8 - CD8+ T cells, cd4 - CD4+ T cells **D.** Dot and whisker plot depicting the LDSC heritability enrichment coefficient from S-LDSC analysis on the Kunkle et al AD GWAS and open chromatin peak sets derived only from the ENCODE monocyte DNase-seq dataset and the Gosselin et al microglia ATAC-seq dataset. P-values for enrichment of each cell type relative to the other are indicated above the dot and whiskers. Whisker ends represent standard errors computed by the S-LDSC software. **E.** Genome browser view of the locus containing the ALPK2 gene which contains a GWAS association with AD by-proxy in the Jansen et al GWAS. Shown are tracks for RefSeq genes, the p-value (−log10 transformed) from the Jansen et al GWAS, and open chromatin peak annotations for CD14+ monocytes (ENCODE), microglia (Gosselin et al), and putamen neurons (Fullard et al). The yellow box indicates the location of the sentinel AD associated variant (rs76726049) at this locus. As visualized, this variant overlaps a monocyte peak but not a microglia peak, suggesting a potential peripheral immune-specific function and not a brain related function for this variant. NOTE: Since this is exploratory analysis to identify potentially relevant cell types, in some cases, we show results from the Jansen et al study and in some cases, we show results from the Kunkle et al study. Overall, if we find a positive hit in one S- LDSC analysis, we include it in future model training since it could be relevant to disease. The remaining S-LDSC analysis plots are provided in the Supplementary Material.

### Monocytes and dendritic cell open chromatin regions are enriched for AD-associated regulatory variants relative to other immune cell types

To compare cell type enrichments within different immune cell types, we reprocessed the single-cell ATAC-seq dataset of immune cell types from Satpathy et al(51) (referred to as “Satpathy immune scATAC” in future). We first grouped cells from the study into 8 major cell types based on the original cluster annotations (monocytes, B cells, CD4+ T cells, CD8+ T cells, dendritic cells, progenitor cells, NK cells, and basophils) to obtain reliable peaks for each of the 8 cell types (Methods) (52). In S-LDSC analysis on this dataset, monocyte and dendritic cell scATAC OCRs displayed a statistically significant enrichment for colocalization with AD risk relative to OCRs for the other immune cell types **(Fig 4C, Fig S3)**.

### Peripheral immune cell types display stronger enrichment for AD-associated regulatory variants compared to brain resident microglia

To investigate the relative roles of brain resident microglia and immune cells in AD, we sought to compare brain resident microglial signals and peripheral immune signals directly in an S-LDSC analysis. To do this, we compared CD14+ monocyte OCRs (DNase-seq from ENCODE: ENCFF231SIM, ENCFF000TBP, ENCFF690LNW, ENCFF000TBL) and microglia OCRs (ATAC-seq from Gosselin et al(49)), Surprisingly, we found that CD14+ monocyte OCRs displayed a significant enrichment relative to microglia OCRs for the Kunkle et al GWAS of diagnosed AD (LDSC coefficient=2.95e-7,p=0.019) **(Fig 4D)**. With the Jansen et al GWAS of diagnosed AD and AD-by-proxy, the CD14+ monocyte datasets displayed a higher enrichment coefficient than the microglial dataset; however, the enrichment was not statistically significant (LDSC coefficient=4.41e-8,p=0.116) **(Fig S3)**. Nevertheless, this suggests that there is a subset of AD associated loci that exert their influence on disease predisposition exclusively through peripheral blood CD14+ monocytes and not through brain resident microglia. For example, a sentinel variant in the Jansen et al GWAS near the ALPK2 locus identified in the Jansen et al GWAS study, rs76726049, lies in a region that is in an OCR in the ENCODE CD14+ monocyte data but not in the Gosselin microglia dataset **(Fig 4E)**, suggesting monocyte-specific regulatory effects for this variant.

### CNN classifiers and regression models trained on open chromatin measurements from AD-relevant cell types can accurately predict regulatory activity from sequence

Having identified peripheral monocytes and microglia as strong candidate cell types for influencing AD predisposition, we trained CNN regression models on the ENCODE CD14+ monocyte and the Gosselin microglia dataset. We conducted model training in a similar fashion to the models trained on the GM12878 DNase-seq dataset (Methods). In addition, we trained a CNN regression model on a putamen neuronal (NeuN+ sorted nuclei) ATAC-seq dataset from Fullard et al(50) (“Fullard NeuN+ ATAC”) since neuron damage is a key consequence of AD. We found that CNN regression models trained on all three of these datasets were able to generalize well and predict open chromatin signals accurately on completely held out chromosome 8 and 9 OCRs in each cell type **(Fig 5A-C, Fig S4)**.

**Fig 5:**
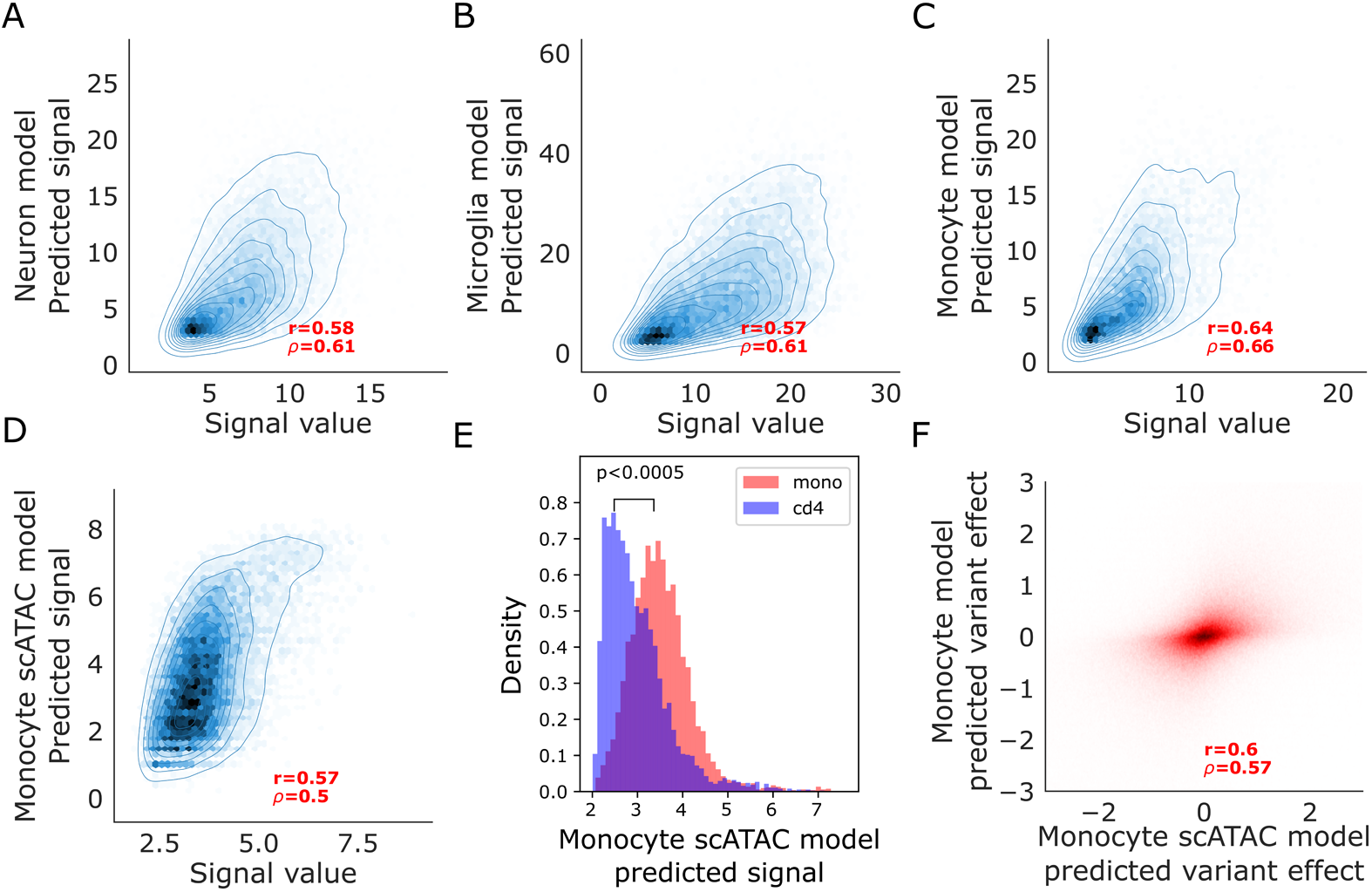
Performance of CNN regression models trained on bulk and single cell ATAC-seq datasets from AD relevant cell types on held out peaks. **A.** Scatter plot of true ATAC-seq signal from the peak caller vs predicted ATAC-seq signal on test set examples (chromosomes 8 and 9) from a CNN regression model trained on open chromatin peaks from the Fullard et al NeuN+ neuronal ATAC-seq dataset **B.** Similar to A. but for a model trained on open chromatin peaks from the Gosselin et al microglia ATAC-seq dataset **C.** Similar to A. and B. but for a model trained on open chromatin peaks from the ENCODE CD14+ monocyte DNase-seq dataset **D.** Scatter plot of true scATAC-seq signal derived from ArchR peak calling on pseudo bulk samples of all monocytes in the scATAC-seq dataset versus predicted signal on test set. (chromsomes 8 and 9) peaks from a CNN regression trained on the monocyte ArchR peaks **E.** Histograms of predicted signal values from a CNN regression model trained on monocyte scATAC-seq peaks from the Satpathy et al dataset on peaks that are exclusively active in monocytes relative to CD4 T cells and peaks that are exclusively active in CD4 T cells relative to monocytes. A Mann-Whitney test was used to test for difference between the means of the predicted signals for the two distributions. p-value from the test is indicated **F. S**catter plot of predicted variant effects from the bulk ENCODE monocyte trained CNN regression model vs the predicted variant effects from the scATAC monocyte trained CNN regression model for all annotated variants in the genome included in the Tanzi variant set

Since we had bulk sorted data for only a couple of immune cell types and we would ideally like to predict the effects of AD variants on multiple immune cell types and identify immune cell type-specific AD variant effects, we also trained models using data from clusters of cells from scATAC-seq, where each cluster’s cells come from a different cell type. Specifically, we applied our CNN regression model training framework on the 8 cell types in the Satpathy et al immune cell scATAC-seq dataset. However, scATAC-seq data presents challenges since there is sparsity in the signal. Sparsity can be introduced by inefficient transposase activity and low cell number for rare cell types which leads to low read coverage in these cell types. Therefore, processing and predicting the signal value in scATAC-seq datasets can be a challenge. To address sparsity and obtain robust signals, we first grouped the original cell clusters from Satpathy et al into 8 major cell classes (monocytes, B cells, CD4+ T cells, CD8+ T cells, dendritic cells, progenitor cells, NK cells, and basophils) and performed peak calling using ArchR (52). Then, we trained 8 different CNN regression models for the 8 cell types modeling ArchR predicted signal at the OCR peak as a function of 500 bp expanded peak summit sequence inputs. Our CNN regression models were able to accurately predict the ArchR signal for all 8 cell type groupings **(Fig 5D, Fig S5)**. Correlation between true signal values and predicted signal values for the holdout test dataset was strong (Pearson’s r between 0.52-0.65 and Spearman’s ρ between 0.49-0.59) for all cell types (**Table 1**), indicative of strong performance.

### scATAC-seq trained CNN models predict cell type-specific signals

Many enhancer variants have effects in only specific cell types and a good model needs to distinguish between enhancers active in a cell type and not in another cell type. Our scATAC monocyte model predicted significantly lower scores for CD4 T cell peaks that are not peaks in monocytes **(Fig 5E)**. Model predictions of variant effect from CNNs trained on bulk CD14+ monocytes and CNNs trained on the scATAC monocytes displayed strong correlations (r=0.6, ρ=0.57) **(Fig 5F).** Combined, this suggests that our scATAC models are able to predict regulatory activity accurately for a cell type as well as capture signals specific to a cell type.

### CNN classifiers and regression models identify candidate causal variants in AD-associated loci that act through specific cell types

To construct an input set of AD associated genetic variants, we identified a set of genetic variants that are in linkage disequilibrium (LD r^2^> 0.5) with sentinel genetic variants identified in two AD GWAS, Jansen *et al.* (9) and Kunkle *et al.* (8). Using this criterion, we constructed a set of 1209 LD-expanded AD associated variants. For scoring a variant, we propagated two sequences through a model, one centered on the reference allele of the variant and the other centered on the alternate allele of the variant. We propagated each of the 1209 sequence pairs through all 11 models and computed predicted allelic skew by computing the difference between the model output for the reference allele carrying sequence from the alternative allele carrying sequence. Since CNN models are known to have difficulties generalizing to areas of the input space which are outside of the training distribution, we focused the initial interpretation of model scores on only genetic variants that overlapped with OCRs in the respective cell type/dataset. To construct a null distribution of predicted variant effect, we made predictions for the full set of common variants that lie in OCRs for a given cell type using the model trained on that cell type. Upon visual inspection, model predictions of variant effect followed a Gaussian-like distribution centered around 0 **(Fig S6)**. Therefore, we used properties of the Gaussian distribution to model variant effect predictions. For each model, we normalized all variant effect scores by dividing them by the standard deviation of allelic skew effect scores for all annotated variants that overlap the respective open chromatin annotations for that model.We did this to mimic a Z-score normalization for a distribution with a mean of 0.From this distribution, we identified outliers by selecting all OCR-overlapping variants that lie at least 2 standard deviations away from the 0 value. We use this cutoff because around 95% of values lie within 2 standard deviations of the mean of a Gaussian distribution. Roughly, this corresponds to a p-value cutoff of 0.05 which is widely used in scientific literature ever since it was described in R.A. Fisher’s tables(58, 59). For additional interpretation of this small set of identified outlier variants, we relaxed our initial overlap criteria and scored their effects using models for other cell types even if the variant did not overlap an OCR in that cell type.

From the initial analysis, we found 7 OCR-overlapping outlier variants that showed predicted effects in CD14+ Monocytes, 3 outlier variants for Microglia, and 1 outlier variant for Neurons. The 7 monocyte outlier variants were identified near the genes MS4A, ABI3, BZRAP-AS1/TSPOAP1, ALPK2, HLA-DRB1, and CLU/PTK2B; the microglia outlier variants were near the genes MS4A, CASS4, and HLA-DRB1; and the neuron variant was near the gene ADAM10. Notably, MS4A, ABI3, PTK2B, and HLA-DRB1 are highly expressed in both microglia (69) and monocytes/macrophages(70–72), and CASS4 is highly expressed in microglia (69). The gene near the neuron outlier variant ADAM10 is a metalloproteinase that forms a major component of α-secretase involved in processing of amyloid β (Aβ) in neurons(73). A complete list of all identified outlier variants can be found in **Table 2**. Of the 7 monocyte candidates, the 4 variants near ALPK2 (rs76726049), CLU/PTK2B (rs28834970), and HLA-DRB1 (rs9270921, rs9271182) did not overlap a microglia peak, suggesting an effect exclusive to peripheral immune monocytes and not brain resident microglia.

Our bulk ENCODE CD14+ monocyte model identified rs28834970 (−3.52 s.d. from 0) near the CLU and PTK2B genes as an outlier. Notably, our scATAC-seq models trained on monocytes (−3.31 s.d. from 0), dendritic cells (−3.2 s.d. from 0) **(Fig S7)**, and basophil models (−3.26 s.d. from 0) all predicted this variant as an outlier, indicating that the effect of this variant may be shared across these peripheral immune cell types. Our models predicted the reference allele (also the risk allele) T for rs28834970 to have an increased regulatory activity in these cell types. This is in line with an earlier study which indicated that the alternative C allele alters a CEBP binding site, which leads to decreased accessibility of the regulatory element and also decreased expression of the nearby gene PTK2B in macrophages (21). Even though the variant did not overlap an OCR in the Gosselin microglia dataset, the model score for this variant was strong (−2.05 s.d. from 0).

Matching the results of S-LDSC analysis, there were several variants where the CNN models predicted a peripheral immune-specific effect on gene regulation. rs76726049 is a sentinel variant identified in the Jansen *et al.* GWAS near the ALPK2 gene **(Fig 6A)**. This variant displayed a large predicted effect (2.3 s.d. from 0) from our ENCODE CD14+ monocyte model. Contrastingly, the variant does not overlap an OCR in the Gosselin microglia dataset (49) and the Gosselin microglia trained model predicted a weak effect for this variant (0.74 s.d. from 0) **(Fig S7)**. In addition, the monocyte classifier predicted a very small non-significant effect for this variant (0.18 s.d. from 0). Combined, this suggests that rs76726049 influences AD predisposition by altering regulatory activity in peripheral monocytes and not in microglia. We conducted model interpretation for rs76726049 sequence pair using DeepSHAP(60) on the monocyte model and we found that the model attends strongly to the index position of the variant for the reference allele carrying sequence but does not do so for the alternate allele carrying sequence **(Fig S8)**. Additionally, the model focuses on adjacent nucleotides on the reference allele carrying sequence suggesting that the surrounding sequence pattern that is important for prediction of monocyte regulatory elements is disrupted by the alternate allele of the sequence. Motif search near the variant revealed Znf341 and Znf530 motifs overlapping these adjacent nucleotides that the model focuses on, suggesting a potential disruption of Znf341 or Znf530 binding by this variant in monocytes. Notably, in differential motif enrichment analysis using Centrimo, both Znf341 (E=1.3e-103) and Znf530 (E=3.2e-407) motifs are enriched in the ENCODE monocyte OCRs relative to the Gosselin microglia OCRs. The identification of the peripheral immune-specific effect of rs76726049 demonstrates the remarkable ability of our models to identify candidate variants influencing AD predisposition and the subtle differences between the very similar regulatory element landscapes of peripheral immune cell types and brain resident microglia.

**Fig 6:**
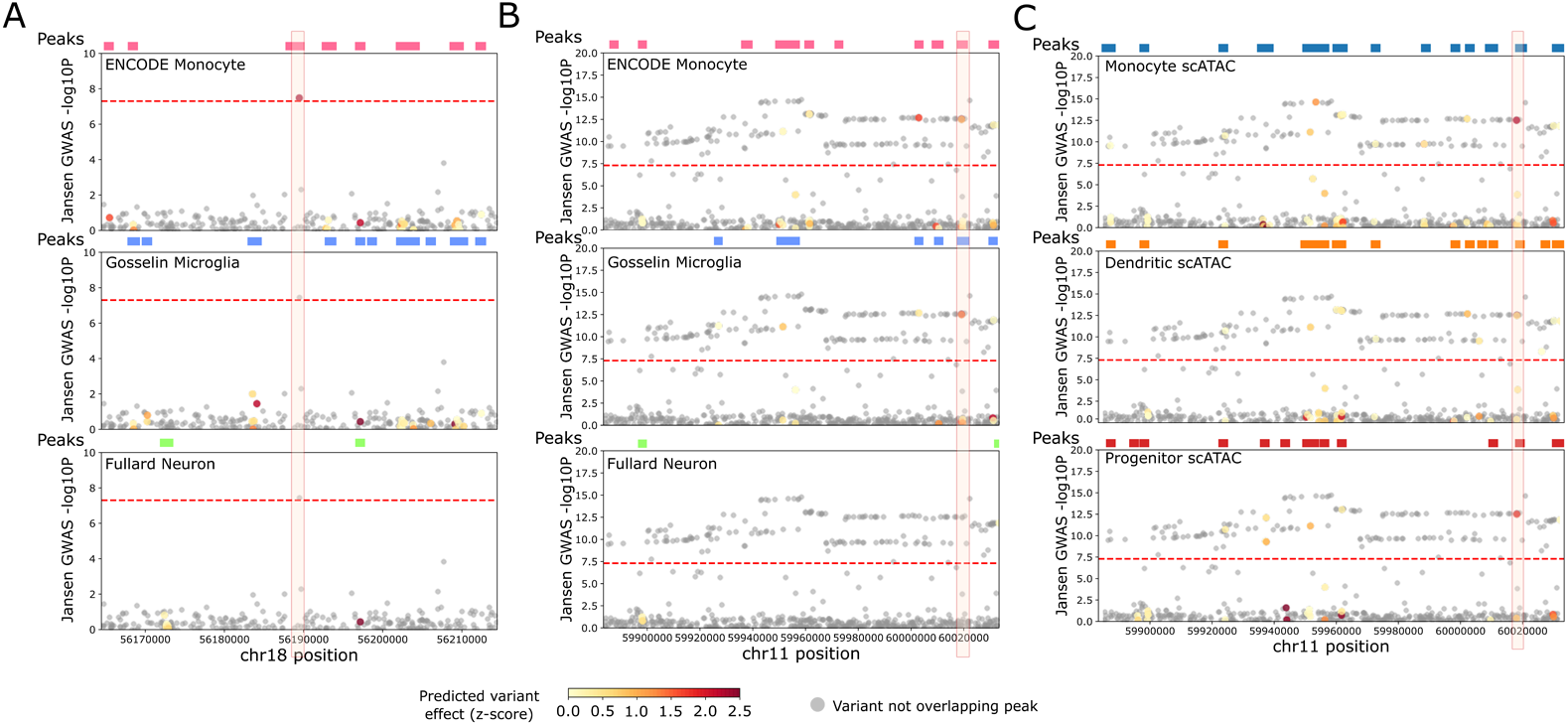
Candidate causal variants identified from models display effects in immune cell types which in some cases, are specific to peripheral immune cell types compared to brain resident microglia. **A.** Genome browser visualization of the AD association region in the Jansen et al GWAS at the locus containing the ALPK2 gene. Tracks include open chromatin annotations from the ENCODE monocyte DNase-seq dataset, the Gosselin et al microglia ATAC-seq dataset, and the Fullard et al NeuN+ ATAC-seq dataset as well as model predictions from the corresponding models trained on that cell type’s dataset. Specifically, the track below each peak annotation track shows the Jansen et al GWAS p-value (−log10 transformed) of each variant in the shown window and the variants are colored by the strength of predicted variant effect from the CNN regression model trained on that corresponding cell type’s dataset. The red highlighted box indicates the location of the sentinel variant at this locus (rs76726049), which overlaps a monocyte peak and is also predicted to have a strong effect on the regulatory activity of that peak from the ENCODE monocyte CNN regression model. Variants that do not overlap open chromatin peaks in the corresponding cell type’s dataset were not scored using the models and are shown in grey. **B.** Similar to A. but for the AD associated region near the MS4A locus. The red highlighted box indicates the location of rs636317 which overlaps both a microglia and a monocyte peak and displays a strong predicted effect from both the monocyte and microglia trained CNN models. **C.** Similar to B. but includes annotations from the scATAC-seq dataset as well as model predictions from the CNN regressions trained on the Satpathy et al scATAC-seq cell types. Shown are scATAC-seq based annotations and predictions for monocyte, dendritic cells, and progenitors but predictions for other cell types were also made. These three cell types displayed peak overlaps with rs636317 in the scATAC-seq dataset and also a significant predicted effect for this variant on regulatory activity of the corresponding overlapping peak.

The monocyte and microglia models both predicted rs636317 near the MS4A locus as an outlier **(Fig 6B)**. This variant has previously been reported to display allelic imbalance on the overlapping ATAC-seq peak in myeloid cells(74) and has been predicted by another CNN model to have a strong effect in microglia(19). Our models predict that the AD risk allele led to lower regulatory activity, and this direction of effect is consistent with the Novikova *et al.* (74) study that showed that the risk allele disrupts a CTCF transcription factor binding site to decrease regulatory activity. rs636317 does not overlap an OCR in the Fullard neuron model, however, it displayed a strong predicted effect **(Fig S7)**, leaving open the possibility that the variant also has an effect in neurons. Our scATAC-seq trained models also called rs636317 as an outlier for monocytes, dendritic cells, and progenitor cell types **(Fig 6C)**. Similar to the bulk models, the scATAC trained models on monocytes, dendritic cells, and progenitors predicted a decrease in regulatory activity for the AD risk allele.

Near the ADAM10 gene, our dendritic cell (3.59 s.d. from 0), natural killer cell (3.01 s.d. from 0), and CD4 T cell (3.53 s.d. from 0) scATAC-seq-trained models identified rs514049 as an outlier. Our bulk neuron model identified a different outlier variant rs395601 near ADAM10 (−2.49 s.d. from 0), which suggests that different variants may contribute to immune-specific and brain-specific effects of ADAM10 gene expression on AD. Notably, ADAM10 displays high expression in peripheral immune cell types (75) as well as neurons, where it is involved in Aβ processing.

## DISCUSSION

Computational genomic approaches have been valuable for linking candidate disease-associated genetic variants to regulatory function in specific tissues and cell types. However, these inferences may be confounded when compared across cell types that are highly similar. Here, we propose a computational framework which enables prediction of variant effects on regulatory activity in disease-relevant cell types. First, we use stratified LD score regression (S-LDSC) to directly compare GWAS association data with open chromatin peaks in multiple cell types. Next, for the implicated cell types, we train regression convolutional neural networks that are able to predict the cell type-specific effect of candidate genetic variants. We are also able to validate the increased sensitivity of our models using MPRA data.

Using this framework, we systematically compare the regulatory impact of genetic variants on multiple neural and immune cell types which have been implicated in the predisposition to AD, some of which are highly similar. Surprisingly, both S-LDSC and the regression CNN approach suggest a stronger role for peripheral monocytes relative to brain-resident microglia. First, while both blood cell types and microglia are enriched for LOAD-associated genetic variants relative to other tissues, we find that blood myeloid cell OCRs display a stronger enrichment than microglia, the brain resident myeloid cell type. This suggests that while many AD associated variants lie in regulatory elements shared between blood and brain immune cell types, a significant subset lie in regulatory elements that are active in only blood immune cell types. Second, several of the candidate FAD-SNPs we identify are predicted by CNN regression models to have a stronger impact on gene regulation in CD14+-monocytes than microglia. While multiple groups have sought to understand the function of regulatory variants in microglia (19, 74), our results indicate both a shared mechanism of AD progression between microglia and blood myeloid cells, and certain exclusive mechanisms of AD progression specifically through blood myeloid cells such as CD14+-monocytes. This necessitates a stronger focus on peripheral blood immune cell types in the future study of AD associated regulatory variant effects.

We acknowledge that the variants unique to CD14+-monocytes may act only in specific microglial states such as activated microglial states associated with AD related inflammation that our microglia datasets might not have captured. Therefore, future research on the function of regulatory AD-associated variants needs to include not only models of inactivated microglia but also other microglial states. Comparing between immune cell types using scATAC-seq data, we find monocytes and dendritic cells are enriched for AD regulatory genetic variants suggesting that a stronger focus on these cell types is also needed in studies of AD regulatory genetic variants. However, this does not support the exclusion of other immune cell types since our previous analysis suggests immune cell types are enriched overall compared to other non-immune tissues and cell types. Further, it is possible that a subset of AD associated variants may act only in stimulated blood myeloid cell states similar to microglia. For example, future studies could include stimulated blood myeloid cell states..

Using our models, we identified both known and novel variants that are predicted to disrupt regulatory activity in the relevant cell types identified in S-LDSC analysis. The most striking example of such variants was rs76726049, a variant near the ALPK2 gene identified as having a CD-14+ monocyte-specific effect relative to microglia from our bulk open chromatin-trained models (**Fig 6**). Model interpretation using DeepSHAP(60) suggested that Znf341 and Znf530 transcription factor binding sites may potentially be disrupted by these variants. Notably, both Znf341 and Znf530 motifs are centrally enriched in the monocyte OCRs relative to the microglia OCRs. In addition, Znf341 and Znf530 are highly expressed in macrophages and monocytes(75) and our analysis suggests that disruption of their binding by rs76726049 could alter monocyte function and influence AD predisposition. Znf341 and Znf530 are highly expressed in immune cell types suggesting a possible role for them in influencing AD through peripheral blood-related processes. More investigation of the ALPK2 locus and the role of these Znf transcription factors is warranted in studies of AD genetic variants that aim to assess their functional effects. More generally, the interplay between these regulatory variants and their effects on ALPK2 expression in influencing AD predisposition needs to be studied. This variant did not overlap a monocyte scATAC-seq peak and did not show a strong effect in the monocyte scATAC-seq trained model. This could be due to low read coverage in monocytes in the scATAC-seq dataset rather than an absence of a regulatory element. Notably, this variant overlapped a scATAC-seq peak identified in the closely clustered dendritic cell population and displayed a moderate predicted effect in the dendritic cell model (1.48 s.d from 0), suggesting that it may not be a false positive based on the bulk CD14+ monocyte model. Since depth of sequencing can lead to low peak numbers in scATAC-seq, this leads to a decrease in the overall number of variants that can be effectively scored by these models. Additionally, artifacts associated with suboptimal clustering can affect downstream analyses. Therefore, our scATAC-seq models may have a high false negative rate. Models trained on deeper and less sparse scATAC-seq datasets will reduce the rate of false negatives. Genetic variants associated with complex diseases can have relatively subtle effects on regulatory activity. Therefore, it is important that computational approaches that aim to predict variant effects can capture these subtle effects. In our framework, we hypothesized that modeling quantitative signals from regulatory assays could improve variant effect prediction performance compared to the commonly trained binary classifiers in the literature. We show that models trained on separate open chromatin measurements (DNase-seq/ATAC-seq) that solely use inherent genetic variation across the genome reference instead of being explicitly fed in sequences with reference and alternate alleles of variants perform comparably well at predicting the direction of variant effects compared to the models trained directly on MPRA data. As hypothesized, we also found that regression models perform better than classifiers at predicting the direction of variant effects in MPRA. Although further hyperparameter tuning of the CNN classifier could potentially bring performance on par with the CNN regression, the minimally tuned regression models are able to predict the subtle differences in regulatory activity induced by many common variants better than classifiers. Therefore, modeling quantitative OCR signals using regression models instead of relying on a binary label which discards continuous information can prove useful in future modeling. In the future, as MPRA datasets become larger and more robust to technical noise, this difference in performance between classifiers and regression models can be better explored.

We rely on MPRA data for validation since it is not influenced by LD as much as other datasets which have been used for variant effect prediction validation such as eQTL data (28, 29, 76, 77), caQTL data, or GWAS catalog membership. To our knowledge, only two studies(65, 66) have validated variant effect predictions from machine learning models directly on MPRA data, which is not confounded by LD. The MPRA-Dragonn study(65) trained CNN models directly on the Sharpr-MPRA dataset(78) in the K562 cell line and validated variant effect predictions on a separate MPRA dataset(79) in K562 that identified 40 expression modulating variants. The authors reported that a model trained on MPRA in the same context was able to predict the direction of effect for 32 of 40 variants correctly. Since the sample size in this MPRA is small, validating model predictions in much larger MPRA datasets like the Tewhey *et al.*(*34*) dataset we used is valuable for demonstrating reliability of model predictions. Hoffman *et al.* (*66*) addressed this issue by successfully validating model predictions in a larger MPRA dataset. As opposed to the commonly trained classifier models in the literature, Hoffman *et al.* used regression to directly model quantitative signals from open chromatin and histone ChIP-seq experiments as a function of the sequence input and were able to show that these models are able to predict MPRA allelic effects reasonably. However, they did not perform rigorous comparisons against classifiers or models trained directly on MPRA data. We conduct a more rigorous extension of this previous study by training models directly on a subset of the MPRA dataset and comparing performance. We also train both classifiers and regression models on open chromatin data and compare performances with these MPRA-trained models.

We show that models trained on DNase-seq perform reasonably well at predicting expression in an MPRA for all sequences. However, these DNase-seq trained models underperform a MPRA-trained model. MPRA datasets are inherently noisy with low replicate concordance and hence, it is possible that this difference in performance may be due to the fact that the MPRA-trained model is overfitting to noise in the MPRA data. In addition, MPRA is primarily an episomal assay whereas DNase-seq and ATAC-seq assay chromosomal DNA directly. Further, open chromatin regions sometimes do not function as enhancers or promoters and this contaminates the training set for open chromatin trained models. All of these factors could be contributing to the drop in performance for both the prediction of expression and variant effect and future work could explore the specific contribution of each factor.

Interestingly, we found that using transfer learning improves performance at predicting expression in MPRA. A model pre-trained on DNase-seq data and fine-tuned on a subset of the MPRA data performed better than models trained solely on DNase-seq or solely on MPRA data. These results suggest that sharing of information and training across assays can help better predict MPRA regulatory activity. For primary tissues, open chromatin datasets are more readily available than MPRA datasets, and training models on open chromatin can help better design MPRA assays. Further, fine-tuning open chromatin pre-trained models on small MPRA datasets can in turn better optimize design of future MPRA experiments. In this way, when there are experimental constraints, it may be possible to attain accurate models of MPRA regulatory activity with a much smaller number of MPRA sequences. This approach could be used to combine open chromatin data and data from a small MPRA to design another small MPRA that would contain likely active regulatory elements, enabling active regulatory elements to be identified without the need for large numbers of cells or especially deep sequencing.

In future work, our model predictions of variant effect in AD can aid in the prioritization of variants for experimental assays or could serve as additional sources of information that help validate experimental results. In the former case, we foresee that variants identified by our models could be prioritized for MPRA experiments in primary microglial or myeloid cells for direct assaying of their regulatory effects. This would be useful when the number of variants/sequences that can be assayed in the experiment is limited due to experimental constraint. Previous MPRAs have largely been conducted in cell lines that are easy to manipulate and allow for assaying of a large number of sequences. Since such cell lines contain chromosomal aberrations and are not especially representative of any primary tissue in a behaving individual, it is important to adapt MPRA experiments to primary tissues. This has proven challenging in the past(80) and machine learning models can help prioritize variants for these assays especially in cases when the number of oligonucleotides that can be assayed is limited. In addition, when large MPRA datasets do become available, our model predictions can serve as auxiliary sources of information that can help validate the measurements of variant effects in future MPRA experiments.

One limitation of our models is that they do not take into account enhancer-gene linking data. In most cases, we use the nearest gene as a proxy for which gene an enhancer may be regulating. However, enhancers are known to loop around to even distant genes. Therefore, some of our candidate variants may actually have an effect on a distal gene. Augmenting our model predictions with better cell type-specific enhancer promoter interactome maps could enable identification of the genes which are affected by our candidate causal variants. Overall, our framework enables cell type targeted studies of variants associated with specific diseases. As opposed to training on hundreds of cell types or tissues(28, 29, 76, 77), we focus on specific cell types/tissues that are identified in S-LDSC analysis to be enriched for disease relevance. We then use individual models targeted on disease relevant cell types/tissues to identify variants influencing AD predisposition as well as their associated regulatory elements and cell types. We rigorously validate our models on experimentally measured variant effects and make our library of models and model predictions available to the community to study the effects of variants associated with AD as well as other immune and neurological disorders. We anticipate that our machine learning approach to identify candidate variants with cell type-specific effects on disease as well as our finding that variants the peripheral immune system may bring about AD will provide the foundation of future investigation of gene regulatory mechanisms involved in the earliest stages of AD.

## DATA AVAILABILITY

Processed open chromatin datasets and Keras model files are available at: http://daphne.compbio.cs.cmu.edu/files/eramamur/ad_variants_resource/ All code for training models, recreating sequence data, generating figures can be found at: https://github.com/pfenninglab/ad_variants https://github.com/pfenninglab/ml-prototypes/tree/master/lcl_regressions

## ACKNOWLEDGEMENTS

We would like to thank Dennis Kostka, Ziv Bar-Joseph, Iliya Lefterov, and Manolis Kellis and members of the Cure Alzheimer’s Consortium for valuable feedback. We thank Dmitry Prokopenko, Rudolph Tanzi, Winston Hide and Lars Bertram for help with variant effect interpretation and for sharing their variant annotation database. We thank Li-Huei Tsai, Gwyneth Welch, and Jemmie Cheng for access to cell type-specific brain H3K27ac profiles. We thank Robert Murphy, Chaitanya Srinivasan, and Andrew Moore for valuable discussions related to transfer learning models. We would like to acknowledge our data sources: the John Stamatoyannopoulos group for generating the GM12878 DNase-seq data, the ENCODE Consortium and the sources of bulk Monocyte open chromatin data, Gosselin et al, Satpathy et al. Lastly, we would like to thank the donors of these datasets.

## CONTRIBUTIONS

ER reprocessed Gosselin et al microglia data, implemented CNN models and trained them on DNase and MPRA, predicted variant effects for bulk models, overall design of manuscript, writing/editing, and compilation. SA performed data preprocessing/peak calling for scATAC dataset, trained scATAC-seq models and predicted variant effects using scATAC trained models, writing/editing, mentored by ER. NT trained transfer learning models and conducted the entire transfer learning analysis, writing/editing, mentored by IMK, input from ER. IMK processed Fullard et al data, preprocessed monocyte data, mentored NT on transfer learning, writing/editing. BNP did data preprocessing and initial exploration of DNase CNN models for MPRA prediction, editing. ARP directed the study, overall design of manuscript, writing/editing.

---

**Fig S1:**
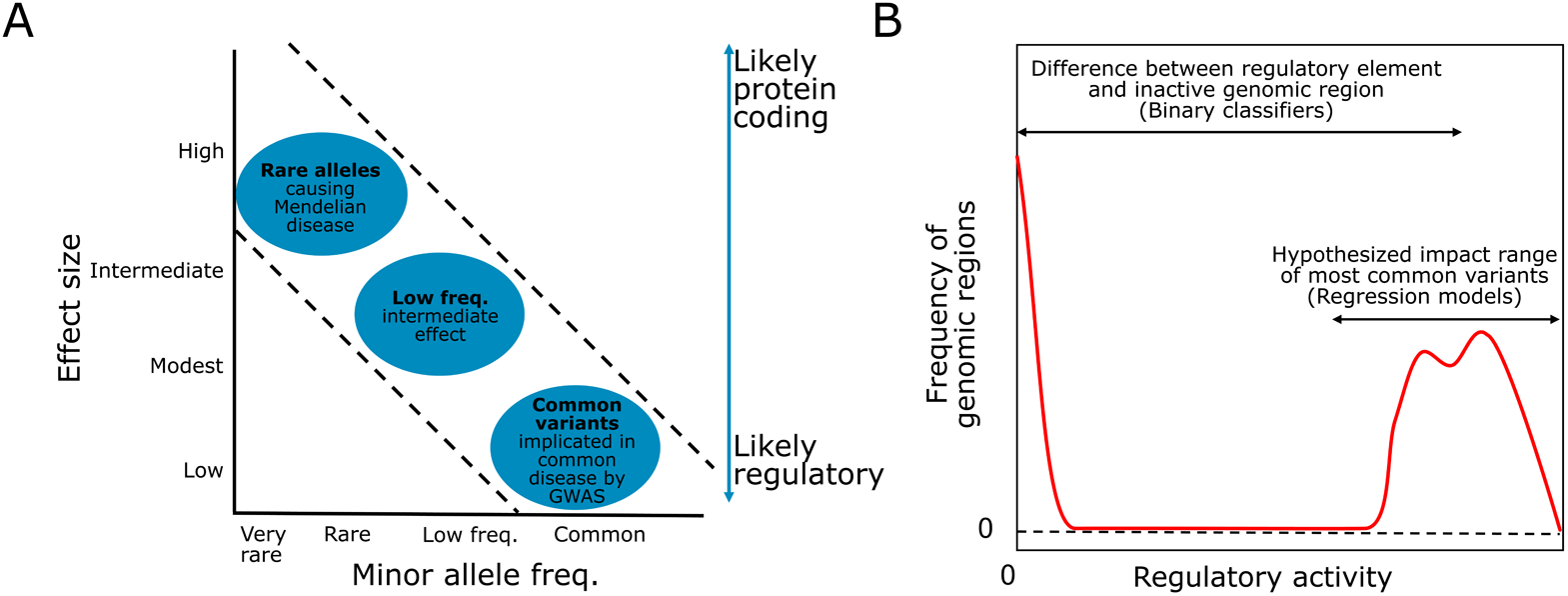
**A.** Cartoon plot showing that common variants implicated in GWAS are likely to have small effects (adapted from Manolio et al 2008) **B.** Cartoon plot showing the distribution of gene regulatory activities for different genomic regions, most genomic regions display 0 or no regulatory activity and a smaller proportion display high regulatory activity. Common variants are likely to have small effects on regulatory activity rather than completely deplete regulatory activity. CNN regressions which model quantitative regulatory signal may be capable of learning these subtle effects of common variants and might help improve upon binary classifiers which put every active genome region into one bin and learn the difference between inactive and active genomic regions

**Fig S2:**
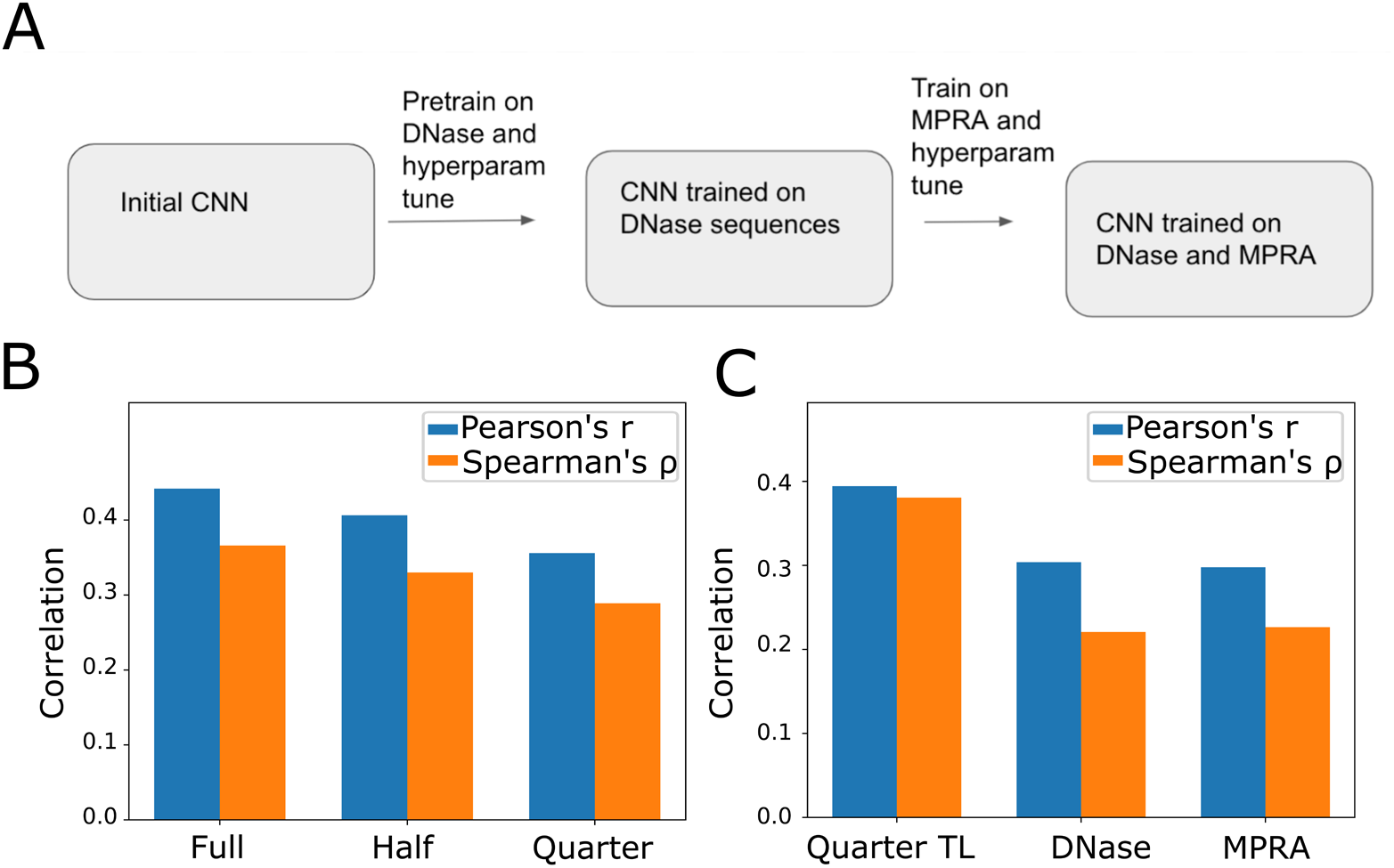
Transfer learning on DNase and MPRA improves prediction of MPRA regulatory activity. **A.** Overview of our transfer learning approach. An initial CNN regression model is pretrained on DNase-seq data and fine tuned on MPRA sequences **B.** Bar chart of Pearson’s and Spearman’s correlation values between true and predicted regulatory activity on sequences in the MPRA test set for transfer learning models fine tuned on different sized subsets (25%, 50%, and 100%) of the MPRA training dataset **C.** Bar chart of Pearson and Spearman correlation values between true and predicted regulatory activity on sequences in the MPRA test set for 1) a transfer learning model fine tuned on quarter (25%) subset of the MPRA training datasets, 2) a non fine-tuned model trained only on DNase-seq, and 3) a model trained on only the quarter subset of the MPRA training datasets

**Fig S3:**
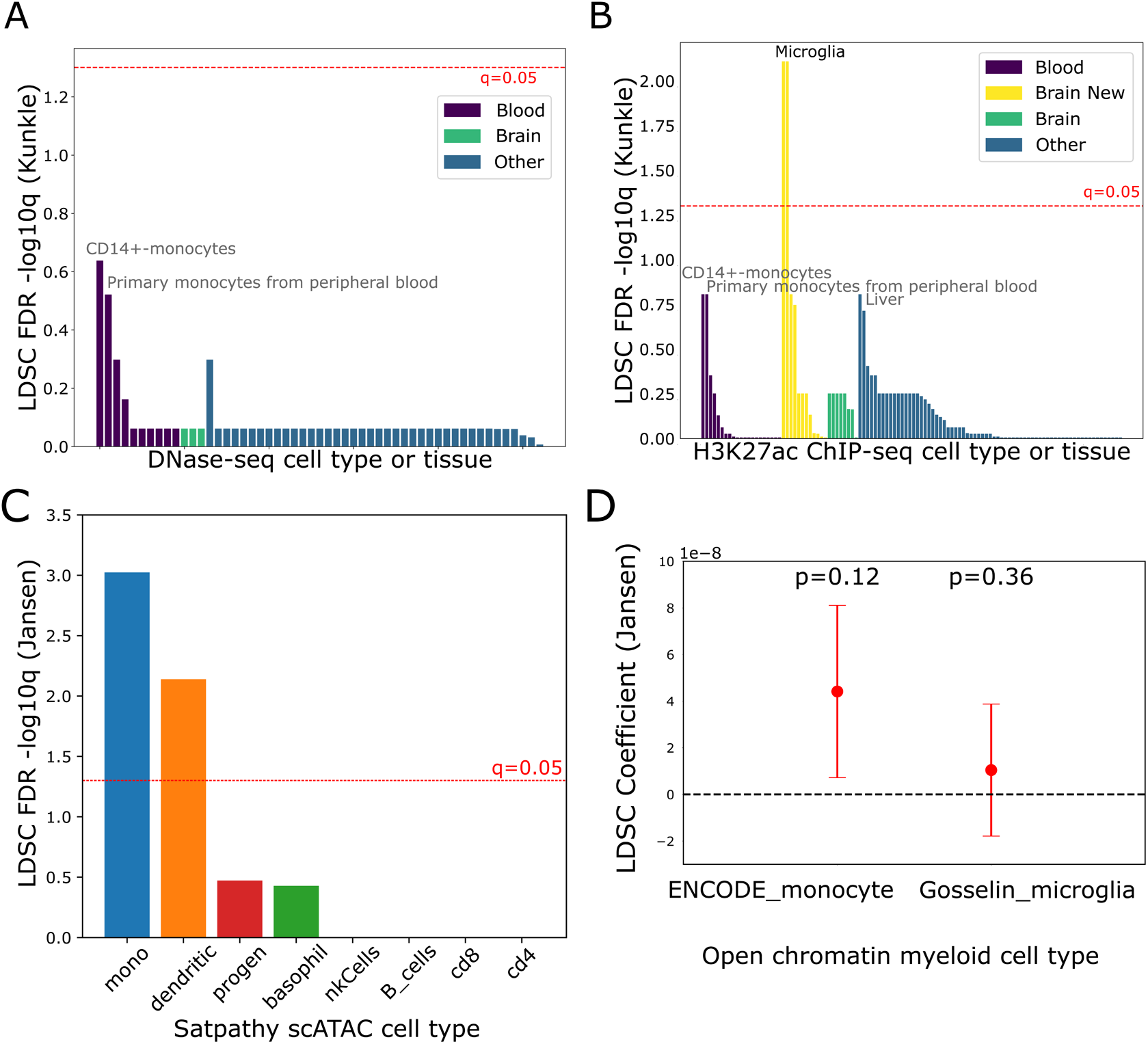
S-LDSC analysis of immune cell types. **A.** Bar chart depicting FDR q-values from an S-LDSC analysis on the Kunkle et al AD GWAS and included DNase-seq peaks from 53 cell types/tissues in the Roadmap Epigenomics dataset. Red line indicates q=0.05, the significance cutoff based on FDR. Tissues are categorized and colored by whether they are derived from brain, blood, or other tissues. **B.** Bar chart depicting FDR q-values from an S-LDSC analysis on the Kunkle et al GWAS and H3K27ac ChIP-seq peaks from 98 cell types/tissues in Roadmap Epigenomics dataset as well as 12 profiles (”Brain New”) from the cell type-specific brain H3K27ac dataset from Ramamurthy, Welch et al. Tissues are categorized and colored by whether they are derived from brain, the new brain dataset, blood, or other tissues **C.** Bar plot showing FDR q-values (−log10 transformed) from S-LDSC analysis on the Jansen et al AD GWAS and open chromatin peaks for 8 immune cell types from the Satpathy et al. scATAC-seq dataset. Red horizontal line indicates q=0.05 which represents the significance level after FDR correction across the 8 tests. **D.** Dot and whisker plot depicting the LDSC heritability enrichment coefficient from S-LDSC analysis on the Kunkle et al AD GWAS and open chromatin peak sets derived only from the ENCODE monocyte DNase-seq dataset and the Gosselin et al microglia ATAC-seq dataset. P-values for enrichment of each cell type relative to the other are indicated above the dot and whiskers. Whisker ends represent standard errors computed by the 5-LDSC software.

**Fig S4:**
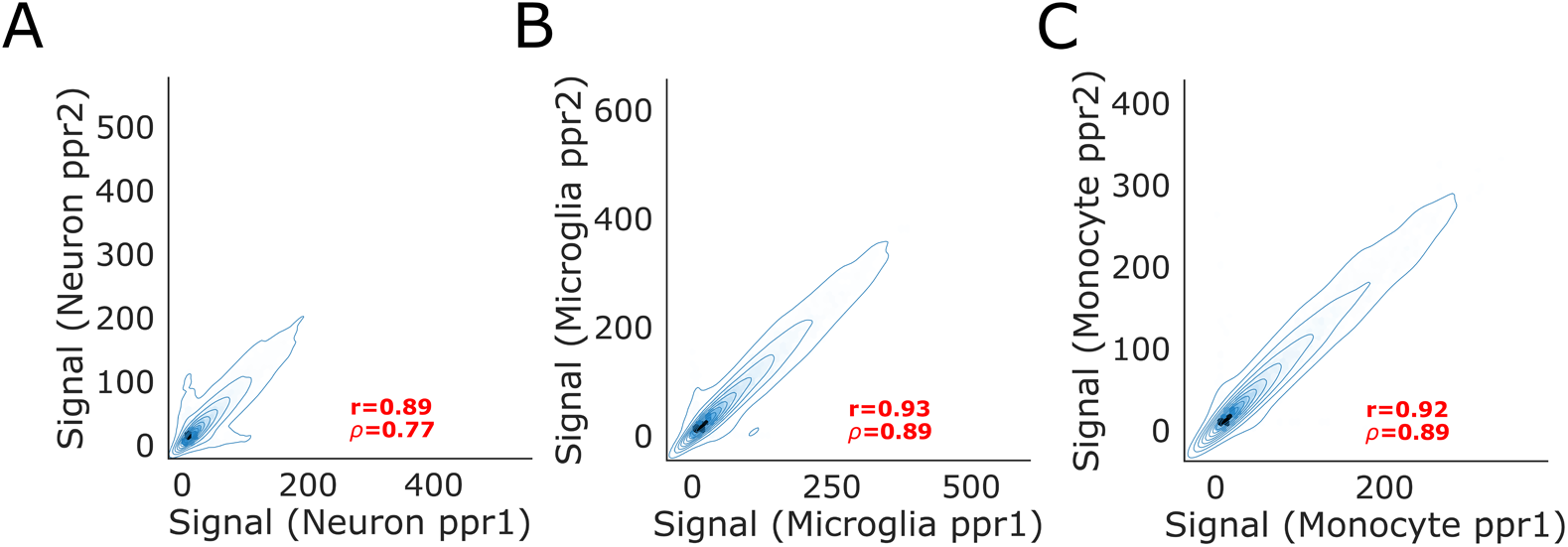
Replicate concordance baselines for the Fullard et al NeuN+ dataset, the Gosselin et al microglia dataset, and the ENCODE monocyte DNase-seq dataset. **A. F.** Scatter plot of true signal values in individual ATAC-seq replicates of the Fullard et al NeuN+ dataset for the held out test set peaks (chromosomes 8 and 9) presented in Fig5A **B.** similar to A. but for the Gosselin et al microglia ATAC-seq dataset and the held out peaks presented in Fig 5B **C.** similar to A. and B. but for the ENCODE monocyte DNase-seq dataset and the held out peaks presented in Fig 5C

**Fig S5:**
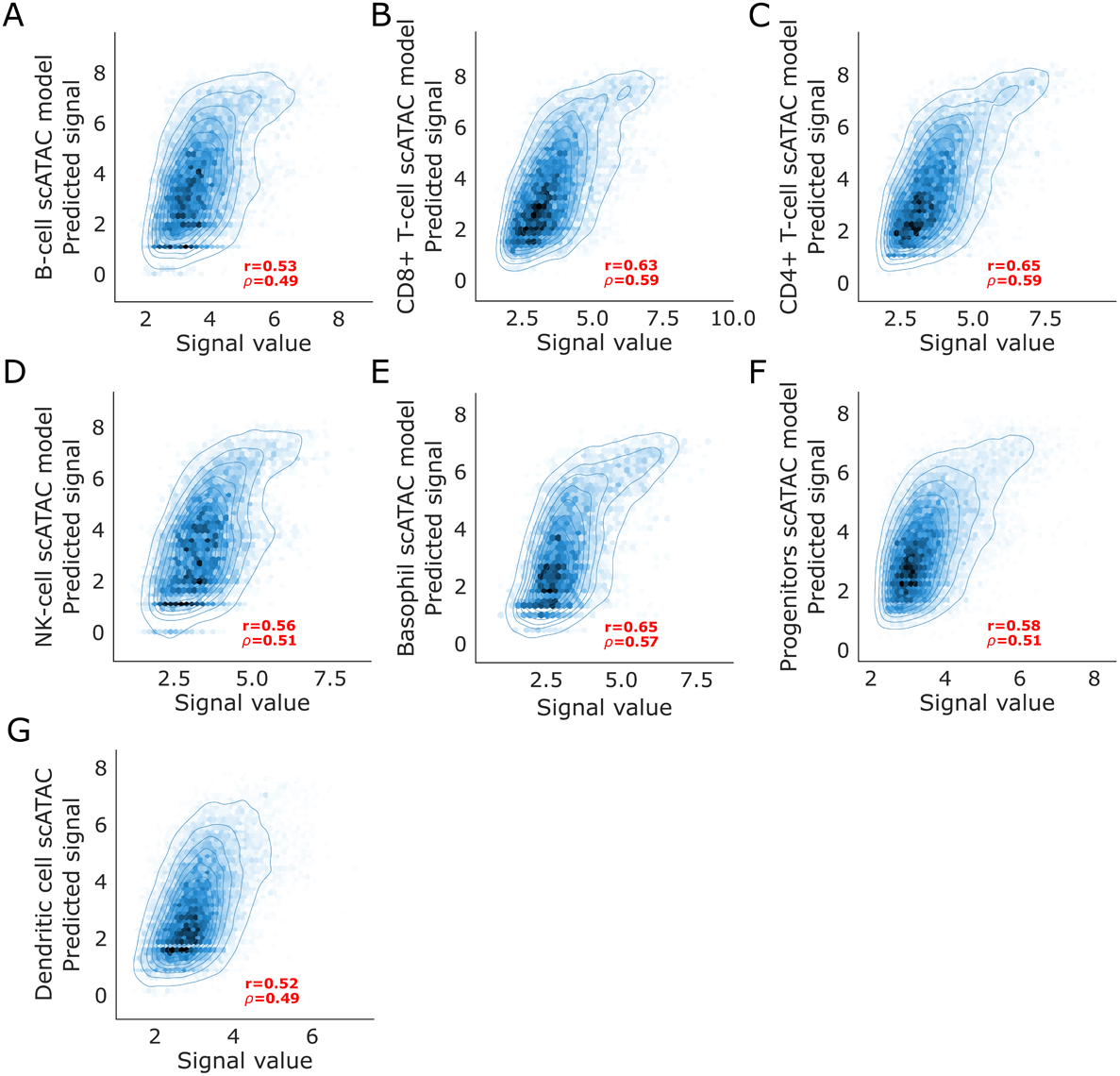
Performance of scATAC-seq trained CNN regression models for remaining immune cell types. Plots show smooth scatter plots of CNN regression model predicted signal vs ArchR called true peak signal for test set peaks (chromosomes 8 and 9) from peaks called on pseudo bulk samples constructed for **A.** B cells **B.** CD8 T cells **C.** CD4 T cells **D.** Natural Killer (NK) cells **E.** Basophils **F.** Progenitors and **G.** Dendritic cells

**Fig S6:**
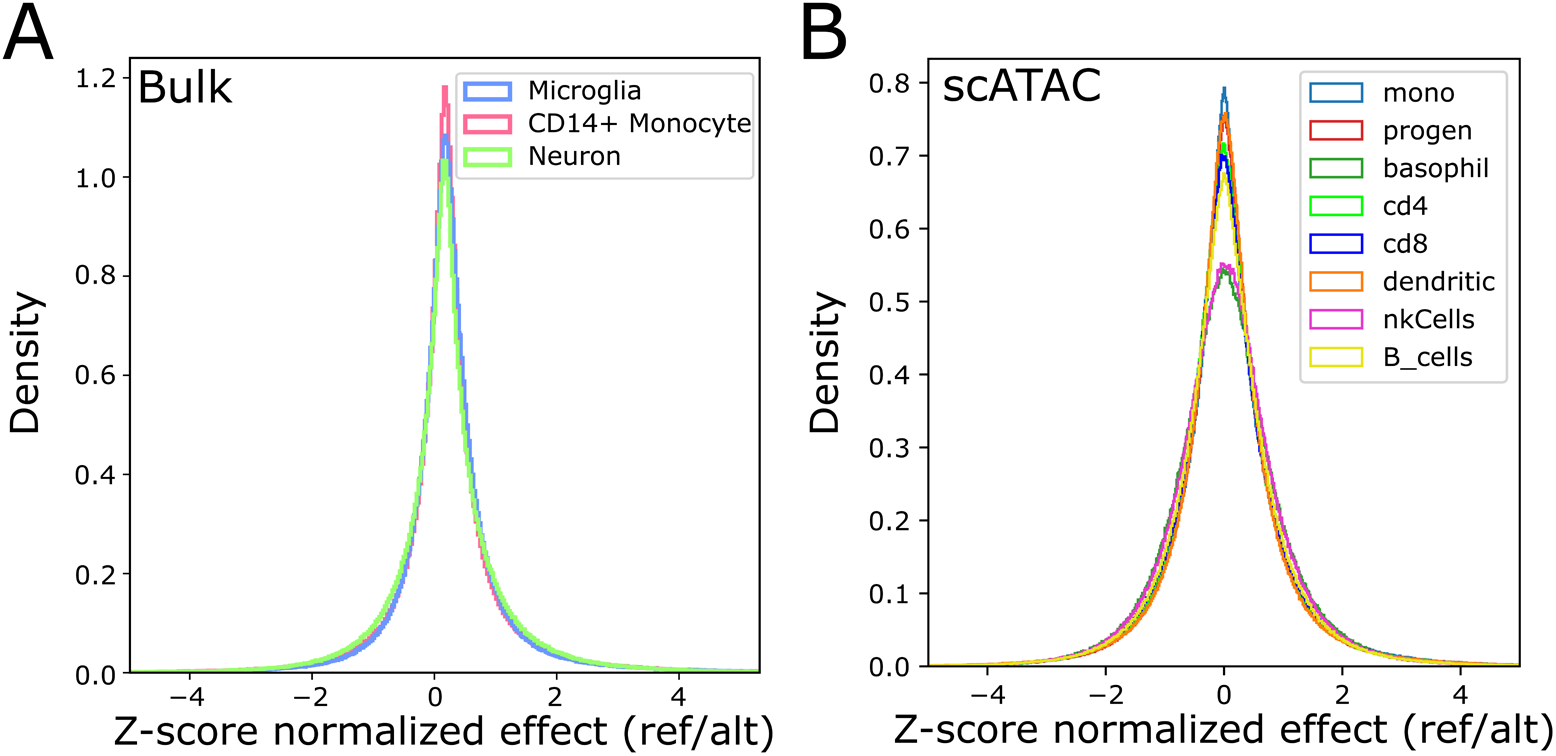
Model predictions of variant effect follow Gaussian-like distributions centered at 0. **A.** Histograms of predicted variant effects for all genomic variants included in the Tanzi variant set from the three CNN regression models trained on the bulk open chromatin datasets (Gosselin et al microglia, ENCODE CD14+ monocyte, Fullard et al Putamen NeuN+) **B.** similar to A. but for CNN regression models trained on the 8 immune cell type peak sets derived from the Satpathy et al scATAC-seq dataset

**Fig S7:**
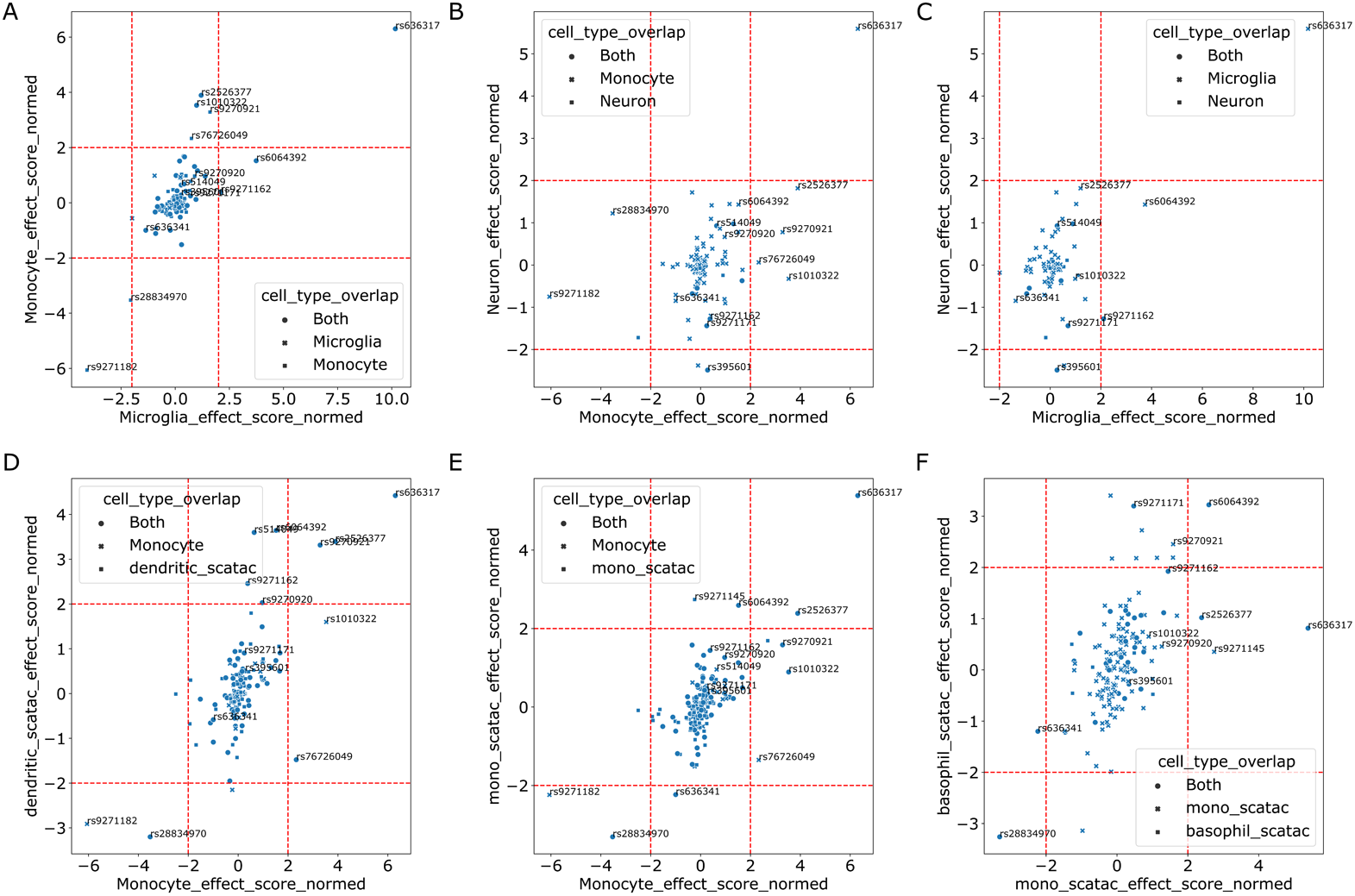
Scatter plots of model predictions of variant effects from two models for AD associated variants that overlap the merged set of OCRs used to train the two models. **A.** Scatter plot of variant effect z-scores from the ENCODE CD14+ monocyte model vs the variant effect z-scores from the Gosselin et al microglia model for variants that overlap peaks in either dataset. The shape of each point indicates whether the variant overlaps OCRs in both datasets or whether the variant only overlaps a monocyte OCR or only overlaps a microglia OCR **B-E** same as A but for model pairs and variants that overlap OCRs in corresponding datasets pairs for other cell types

**Fig S8:**
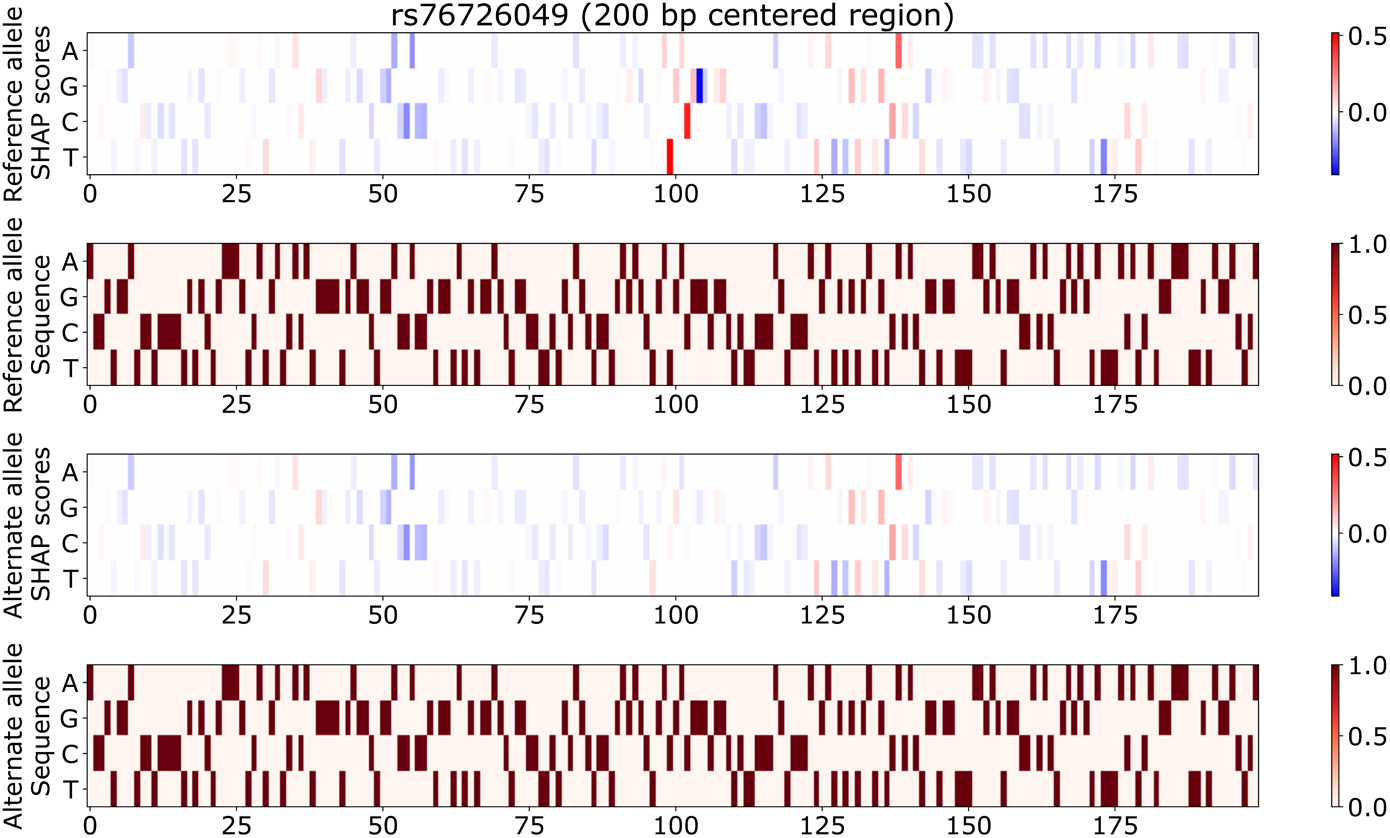
DeepSHAP contribution scores show that the monocyte model focuses on the reference allele as well as adjacent nucleotides on the reference allele carrying sequence for rs7626049. From top to bottom. DeepSHAP contribution scores for the middle 200 bp of for the 1000 bp reference allele carrying sequence. One hot encoding of the middle 200bp of the reference allele carrying sequence. DeepSHAP contribution scores for the middle 200 bp of for the 1000 bp alternate allele carrying sequence. One hot encoding of the middle 200bp of the alternate allele carrying sequence.

**Fig S9:**
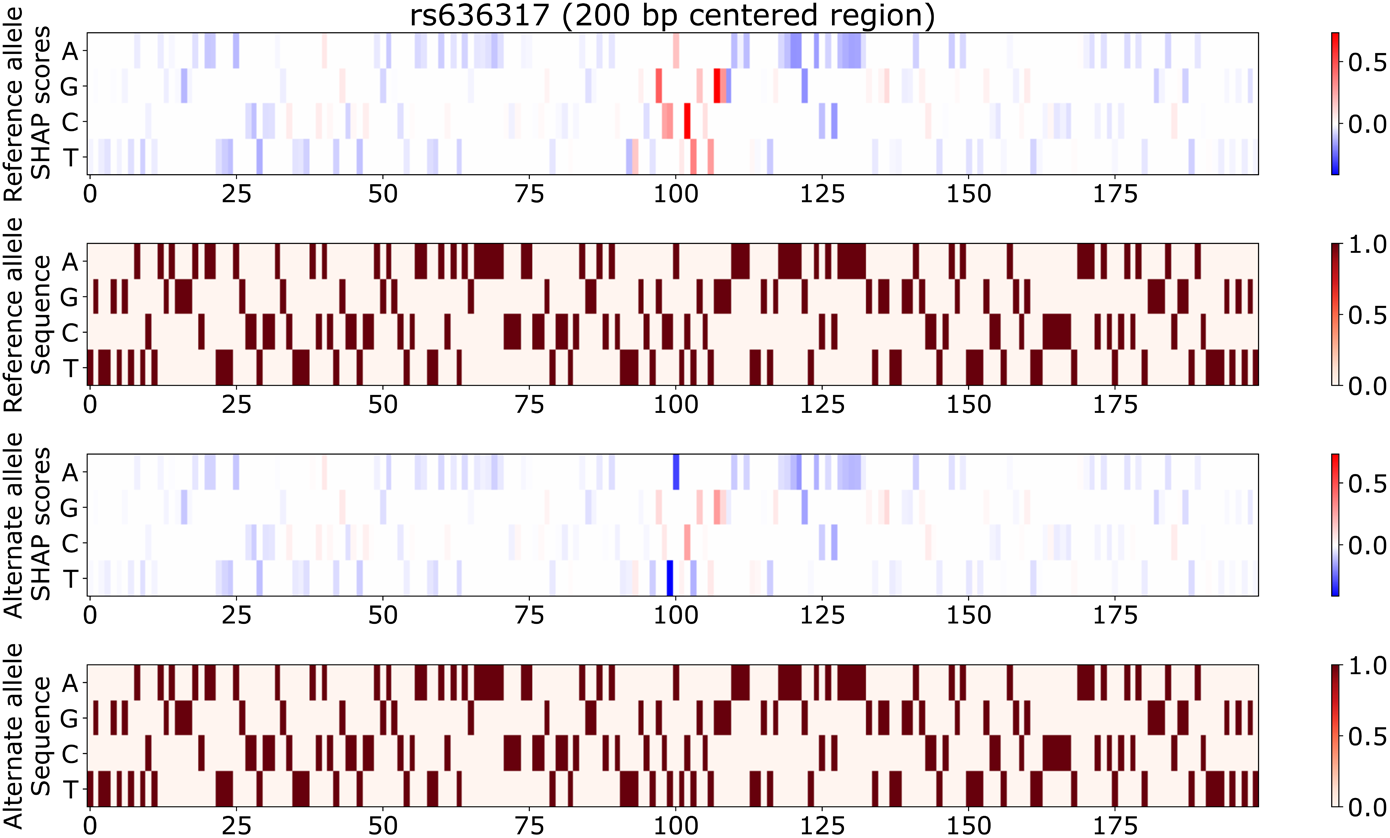
DeepSHAP contribution scores show that the monocyte model assigns positive contribution scores to the reference allele and surrounding nucleotides on the reference allele carrying sequence and negative contribution scores to the alternate allele as well as adjacent nucleotides on the alternate allele carrying sequence for rs636317. From top to bottom. DeepSHAP contribution scores for the middle 200 bp of for the 1000 bp reference allele carrying sequence. One hot encoding of the middle 200bp of the reference allele carrying sequence. DeepSHAP contribution scores for the middle 200 bp of for the 1000 bp alternate allele carrying sequence. One hot encoding of the middle 200bp of the alternate allele carrying sequence.

